# Differential methanogenic archaea-induced TLR8-dependent signaling is governed by NF-κB p65- and STAT1/2-controlled gene classes

**DOI:** 10.1101/2025.09.30.679018

**Authors:** Fan Xu, Johann Sieverding, Peiling Zhou, Shoshant Sah, Viktoria Weinberger, Christine Moissl-Eichinger, Inken Wohlers, Holger Heine

## Abstract

Prevalent in the human gut yet lacking canonical cell wall-derived host-recognition motifs found in bacteria, human-associated archaea *Methanosphaera stadtmanae* and *Methanobrevibacter smithii* show immune activity and disease links but remain largely underexplored. Time-resolved RNA-seq in human PBMCs identified an immune program shared with bacterial or viral stimuli across both archaeal species, yet their kinetics diverged, with *M. stadtmanae* inducing earlier and stronger activation and *M. smithii* eliciting more gradual responses. Among conserved early-upregulated genes, two classes emerged: Class I was preferentially induced by *M. stadtmanae*, whereas Class II was similarly induced by both species. To investigate the origin of the Class I/II phenotype, we measured uptake and applied TLR8 inhibition, finding greater early uptake with *M. stadtmanae* and broad TLR8 dependence across readouts. Consistent with this gene-class separation, ChIP-qPCR showed that *M. stadtmanae*, but not *M. smithii*, strongly induced NF-κB p65 binding at representative Class I promoters, while STAT1/2 binding at representative Class II promoters occurred with both stimuli. Dose-response analyses with RNA inputs established distinct activation thresholds, with Class II at low dose (via STAT1/2) and Class I only at higher dose (via p65). Together, these findings support an input-tuned logic in which archaeal inputs and RNA sensing gate TLR8-dependent immune programs.

## Introduction

Archaea, long regarded as extremophiles inhabiting environments such as hydrothermal vents and hypersaline lakes^1, 2^, are increasingly recognized to encompass mesophilic species widely distributed in temperate ecosystems, including soil, freshwater, and marine habitats^3–5^. They are also stable members of human-associated microbial communities on the skin^6–8^, in the oral cavity^7, 9–12^, and within the respiratory^7, 8, 13, 14^ and digestive tracts^8, 15–19^. Lifestyle factors such as diet and urbanization are also associated with variation in the gut archaeome^20^.

In the human gut, methanogenic archaea thrive under anoxic conditions by consuming bacterial fermentation-derived CO_2_ and hydrogen, thereby contributing to cross-feeding interactions and microbiome stability^16, 21, 22^. Recent multi-cohort analyses further support this view by showing that gut methanogens are embedded in structured archaea-bacteria co-occurrence networks, including associations with fiber-degrading^23^ and butyrate-producing bacterial taxa^24^. Despite this ecological relevance, the human archaeome remained under-characterized for years because standard 16S rRNA gene-based approaches often under-detect archaea, but archaeal-targeted primers^7, 25^ and metagenomic sequencing and genome binning have considerably expanded the catalog of human-associated archaea^17^.

Although minor in diversity, methane-forming archaea can constitute up to 4% of the gut microbiome, underscoring their ecological relevance^26^. As currently understood, archaeal taxa in the human gut are dominated by the order *Methanobacteriales*, mainly *Methanobrevibacter smithii*, *Methanobrevibacter intestini*^27^ and *Methanosphaera stadtmanae*^28^. Methanogenic archaea have been associated with overall health^29, 30^ and longevity^31^. *Methanobrevibacter* local overabundance has been linked with periodontitis^9, 10, 12^, constipation^32^, Parkinson’s disease^33^, and colorectal cancer^34^, whereas depletion has been reported in inflammatory bowel disease (IBD)^35, 36^. Likewise, *M. stadtmanae* exhibits pro-inflammatory properties in IBD^15^, but its early-life colonization is linked to reduced childhood asthma risk^37^. These apparently conflicting associations suggest that the effects of methanogenic archaea are context dependent and likely shaped by abundance, strain composition, community background, diet, and host immune tone.

The gut microbiota profoundly influences immune homeostasis through its metabolic outputs, such as short chain fatty acids, and via direct microbial-immune interactions that shape host immune responses^38–41^. While bacterial commensals are well-established immunomodulators, the mechanisms by which archaea interact with the immune system remain poorly understood, in part because archaea appear to lack canonical cell wall-derived immunostimulatory molecules. In host defense, bacteria are sensed through diverse pathogen-associated molecular patterns (PAMPs), including LPS by TLR4, and lipopeptides by TLR2^42^, peptidoglycan by NOD1 and NOD2^43^, unmethylated CpG DNA by TLR9^44^, and bacterial RNA by TLR7/8^45^. Viruses are mostly sensed through their nucleic acids, although viral proteins may also contribute to innate immune recognition^46^, and viral RNA is detected by endosomal TLR7/8 or cytosolic sensors after entry^47^. Archaea occupy a unique position between these paradigms; they lack canonical bacterial PAMPs such as LPS or classical peptidoglycan^48^. Our previous work, along with that of others, has highlighted the immunogenicity of *M. smithii* and *M. stadtmanae* in human peripheral blood mononuclear cells (PBMCs) and monocyte-derived dendritic cells (moDCs), and provoke pulmonary immune activation in murine models, typified by tumor necrosis factor-alpha (TNF) and interleukin-1 beta (IL-1β) secretion^15, 49–51^. Our work demonstrated that archaeal immune recognition is primarily driven by RNA^51^. Cytokine induction by *M. stadtmanae* strictly depends on phagocytosis and endosomal acidification, and knockout or pharmacologic inhibition of TLR8 nearly abolished the response, establishing archaeal RNA as the dominant trigger of innate sensing. Interestingly, prior study from us showed that dendritic-cell recognition of the commensal *Lactococcus lactis* G121 and its RNA via endosomal TLR8 is required for its allergy-protective effects^52^, highlighting TLR8 as a promising axis for therapeutic modulation. Indeed, several selective TLR8 agonists (e.g., motolimod, selgantolimod, DN052) are in clinical evaluation for oncology and chronic hepatitis B and are being explored as vaccine adjuvants^53–55^.

More recently, additional routes of immune engagement have been uncovered. Archaeal glycerolipids can activate the C-type lectin receptor Mincle^56^, and extracellular vesicles (EVs) released by *M. smithii* and *M. stadtmanae* have been shown to deliver immunomodulatory cargo, although their precise active components remain undefined^57^. Despite these advances, most studies to date have examined only a narrow set of cytokine readouts and have not addressed broader transcriptional programs and regulatory circuits engaged by methanogenic archaea. A system-level approach would not only allow a more comprehensive comparison with bacteria and viruses in terms of immune recognition but could also help explain the nature of archaea in disease associations. In addition, comparative analyses are particularly lacking for the two important gut methanogens, *M. stadtmanae* and *M. smithii*, which inhabit the same host environment yet differ in fundamental biology that may shape host interaction.

*M. stadtmanae* exhibits a highly restricted methanogenic metabolism and a relatively compact genome, whereas *M. smithii* is a dominant gut methanogen and displays genomic and metabolic adaptations to the intestinal habitat^58^. Nevertheless, comparative genomic analyses suggest that both species are adapted to a host-associated intestinal lifestyle, as each encodes a distinct repertoire of adhesin-like surface proteins that differs in composition and domain architecture^59^. In addition, the two species also differ in their immunostimulatory capacity, with *M. stadtmanae* generally eliciting stronger pro-inflammatory responses than *M. smithii* in both the human and murine system^15, 49, 50^. This gap hinders mechanistic insight into how conserved versus species-specific pathways are programmed in human immune cells.

Here we present a genome-wide transcriptomic analysis of human PBMCs, and PBMC-derived monocytes stimulated with the important gut methanogens, *M. stadtmanae* and *M. smithii*, leveraging time-resolved transcriptomics to resolve the dynamics of host responses. We advance a unifying model in which cellular entry routes and TLR8-mediated sensing of archaeal RNA drive distinct transcriptional programs, thereby offering mechanistic insight into archaeal-host crosstalk, and providing a conceptual reference to innate programs previously characterized for bacterial and viral stimuli, while motivating future investigations into input-tuned innate signaling.

## Materials and Methods

### Ethics Approval Statement

Approval for these studies was granted by the Institutional Ethics Committee of the University of Lübeck (Lübeck, Germany; Protocol No. Az. 12-202A) in accordance with the Declaration of Helsinki. Written informed consent was obtained from healthy adult donors. Healthy adult donors were included without selection based on sex. Where available, sex information was based on donor information collected at recruitment. As the study was not designed or powered to assess sex-specific effects, no stratified analyses were performed.

### Archaea Growth and Media

The human gut-derived strains *Methanobrevibacter smithii* (DSM 2375, strain ALI) and *Methanosphaera stadtmanae* (DSM 3091, type strain) were obtained from the German Collection of Microorganisms and Cell Cultures (DSMZ) GmbH (Braunschweig, Germany). For the cultivation of all methanogens standard MS medium was used as previously describe^27^. For immune cell stimulation experiments, exponentially growing *M. stadtmanae* and *M. smithii* cells were harvested by centrifugation at 3200× g for 30 minutes, washed, and resuspended in aerobic 50 mM Tris-HCl (pH 7.0).

### Isolation of Archaeal RNA and DNA

For nucleic acid isolation, *M. stadtmanae* and *M. smithii* cells were cultured as previously described, harvested at 4°C by centrifugation at 3200× *g* for 30 minutes, and lysed in liquid nitrogen using a Mikro-Dismembrator S laboratory ball mill (Sartorius) for 3 minutes at 1600 bpm. RNA was extracted using the TRIzol method, followed by DNase I treatment. For isolation of extracellular archaeal RNA from culture supernatants, cell-free supernatants were generated by centrifugation (3200× g for 30 minutes) to pellet intact archaeal cells and residual particles, followed by passage through a 0.22 µm filter. DNA was isolated using the Wizard Genomic DNA Purification Kit (Promega) according to the manufacturer’s protocol. The concentration and purity of nucleic acids were assessed using a DS-11 spectrophotometer (DeNovix). RNA with an A260/A280 ratio of ≥2.0 was considered pure; DNA with an A_260_/A_280_ ratio of approximately 1.8 was considered as pure DNA.

### Cell Culture

PBMCs were isolated from heparinized donor blood of healthy adult donors using gradient centrifugation with Biocoll (Merck)^60^. Monocytes were then purified using the MagniSort™ Human CD14 Positive Selection Kit (BioLegend, #480048)^61^. Monocyte-derived dendritic cells (moDCs) were subsequently generated by culturing CD14^+^ monocytes in complete medium supplemented with GM-CSF (50 ng/mL) and IL-4 (1000 U/mL) for 7 days, as described previously^62^. On day 3 and day 5, cultures were refreshed by removing half of the spent medium and replacing it with an equal volume of fresh complete medium containing GM-CSF and IL-4 at 2× concentration to maintain the intended final cytokine concentrations. All cells were cultured in RPMI 1640 medium with stable glutamine, supplemented with 10% FCS and antibiotics 100 U/mL penicillin and 100 µg/mL streptomycin (P/S; all from Merck)], hereafter referred to as complete medium. Cultures were maintained in a humidified incubator at 37°C with 5% carbon dioxide. BLaER1 cells were kindly provided by Thomas Graf (Center for Genomic Regulation, Barcelona, Spain). Cells were cultured under the same conditions as moDCs and maintained at a density of 1 × 10^5^ to 2 × 10^6^ cells/mL. For transdifferentiation, cells were seeded at 3 × 10^5^ cells/mL and cultured for 7 days in complete medium supplemented with 10 ng/mL IL-3 (PeproTech, # 200-03), 10 ng/mL M-CSF (PeproTech, # 300-25), and 100 nM β-estradiol (Sigma-Aldrich, #E2758), as previously described^63^. For stimulation experiments, transdifferentiated cells were replaced in complete medium without IL-3, M-CSF, or β-estradiol.

### Cell Stimulation

For the ELISA experiment, PBMCs, monocytes, and moDCs were seeded at a density of 5×10^4^ cells per well in 96-well flat-bottom plates (200 µL total volume) and stimulated for 4 or 24 h at 37°C under 5% CO_2_. For the RT-qPCR experiment, PBMCs and monocytes were plated at 1.5×10 cells per well in 6-well flat-bottom plates (2 mL total volume) and stimulated for 1, 2, 4, 6 or 24 h at 37°C under 5% CO_2_. The whole cells of *M. stadtmanae* and *M. smithii* were applied at the concentration of 10^7^ cells/ml unless otherwise specified. The chemically synthesized TLR8 compound ligand TL8-506 (InvivoGen, #tlrl-tl8506) was used at a concentration of 1 µg/mL. Archaeal RNA was pre-complexed with the liposomal transfection reagent DOTAP (Sigma-Aldrich, #11202375001) at a ratio of 10 µL DOTAP per 1 µg RNA or DNA in 50 µL pure RPMI 1640. The DOTAP/RNA or DNA complex mixture was incubated for 10 min at room temperature to allow complex formation before stimulation. Total RNA or DNA was added at the indicated concentration. Under our standard stimulation conditions, this procedure resulted in a final DOTAP concentration of 10 µg/mL. For inhibition experiments, 5 µM of the TLR8 inhibitor CU-CPT9a (InvivoGen, #inh-cc9a) or 2 µM cytochalasin D (Sigma-Aldrich, #C2618) was added to the cells 1 h at 37°C prior to stimulation with archaea or TL8-506. As both CU-CPT9a and cytochalasin D were dissolved in DMSO, the corresponding control groups received the same volume of DMSO.

### Cytokine Measurements

Cytokine concentrations in the supernatants were measured after 24 h or 4 h stimulation using commercial ELISA kits specific for IL-1β, IL-6, TNF-α, and CCL2 (Thermo Fisher).

### Multiplex immunoassay platform

A multiplex analysis was conducted at a central laboratory using banked supernatants and the 20-cytokine U-PLEX™ immunoassays (Meso Scale Discovery)^64^, which include multiple cytokines associated with immune responses to viral infections. The assays followed the manufacturer’s protocol and were analyzed using the MESO QuickPlex SQ 120 imager (Meso Scale Discovery). The measured analytes included granulocyte-macrophage colony-stimulating factor (GM-CSF), interferon (IFN)-γ, IFN-β, IFN-α2a, interleukin (IL)-1 receptor antagonist (RA), IL-1β, IL-4, IL-5, IL-6, IL-7, IL-8, IL-9, IL-10, IL-12p40/IL-12B, C-C motif ligand 2 (CCL2), also known as monocyte chemotactic protein 1 (MCP-1), CCL3, IFN-γ-inducible protein 10 (IP-10), also referred to as C-X-C motif chemokine ligand 10 (CXCL10), tumor necrosis factor (TNF)-α, and vascular endothelial growth factor (VEGF)-A. Cytokine concentrations in the supernatants were measured after 24 h or 4 h stimulation.

### RNA-Seq and data processing

RNA extraction was performed once all samples analyzed in this work were collected and stored at -80°C using the innuPREP RNA mini Kit (IST). Subsequent steps included RNA quality control, purification of poly-A RNA, RNA fragmentation, strand-specific, random-primed cDNA library preparation, and generation of 150 bp paired-end sequencing data on an Illumina® NovaSeq 6000 platform. Two RNA-Seq experiments were performed (i) to characterize host transcriptional responses to methanogenic archaea and (ii) to assess the role of TLR8 signaling, respectively. In the first experiment, PBMCs from nine healthy donors were stimulated with either *M. smithii* or *M. stadtmanae* for 4 or 24 h, or left unstimulated for 24 h as a control, resulting in five experimental conditions. To ensure temporal consistency and minimize batch or circadian effects, all samples were harvested simultaneously, with 4 h stimulations initiated accordingly. Each condition included nine biological replicates, totaling 45 samples. The second experiment focused on the impact of TLR8 inhibition. CD14^+^ monocytes isolated from PBMCs of three donors were stimulated for 4 h with *M. smithii* or *M. stadtmanae*, with or without a TLR8 inhibitor. Together with unstimulated and inhibitor-only controls, this resulted in six experimental conditions and 18 RNA-Seq samples.

The RNA-Seq analysis was conducted using a community-developed pipeline from the Snakemake workflow catalog (https://github.com/snakemake-workflows/rna-seq-kallisto-sleuth) (version 2.8.4). First, reads were trimmed and transcript abundance estimated with Kallisto^65^. Then, differential expression analysis was performed with Sleuth^66^ using a joint model across all conditions, with the donor included as a covariate in the statistical model. Multiple testing correction was performed within Sleuth using the Benjamini-Hochberg method, and adjusted P values reported in the manuscript correspond to FDR-adjusted q values. Quality control of sequencing reads was performed with FASTQC (https://www.bioinformatics.babraham.ac.uk/projects/fastqc), and an aggregated quality report, including alignment metrics, was created using MultiQC^67^. Transcript annotations are based on GRCh38 and Ensembl release 114. Functional enrichment analysis was available in the workflow and uses SPIA^68^ for computing enriched KEGG pathways^69^. Significant differences across all five or six conditions are computed using a likelihood ratio test. Significance of the effect estimates of each condition versus the respective control condition is tested using a Wald test. For expression profiling, the most highly expressed transcript of each gene was kept, and transcripts were sorted by expression variance across samples. Hierarchical clustering was performed using 1 minus Pearson correlation as the distance metric, based on the top 100 most variable genes (across samples). Principal component analyses were performed using the 1000 most variable transcripts. Some gene symbols were represented more than once in the Sleuth output as they were assigned to multiple Ensembl gene IDs. To obtain a non-redundant results table for downstream reporting and visualization, duplicated gene symbols collapsed by retaining the entry with the highest expression level.

### Gene Ontology enrichment analysis

Gene Ontology (GO) enrichment analysis was performed using the DAVID Bioinformatics Resources 6.8 (https://david.ncifcrf.gov/) to explore the functional categories enriched among up/down-regulated genes. Analyses were conducted for the three GO categories: Biological Process (BP), Molecular Function (MF), and Cellular Component (CC). Enrichment results were filtered using the following criteria: P-value < 0.05 and gene count > 2. The top five or ten most significantly enriched terms from each GO category were selected for visualization.

### Protein-protein interaction (PPI) network analysis

Protein-protein interactions among the 443 core genes were retrieved from the STRING database (version 12.0; accessed via the web interface) using *Homo sapiens* as the reference organism and applying a high-confidence interaction score threshold (≥ 0.9). Both experimentally validated and database-curated interactions were included. The resulting interaction table was imported into Cytoscape (version 3.10.3) for network visualization. Node size and node color were both scaled according to node degree (number of connected edges), with larger and more intensely colored nodes indicating higher connectivity.

### Reactome Pathway Enrichment Analysis

Pathway enrichment analysis was performed using the Reactome Pathway Browser (https://reactome.org). Differentially expressed genes were uploaded to the web tool, and over-representation analysis was conducted against the Reactome human pathway database. Significance was assessed using the hypergeometric test with Benjamini-Hochberg false discovery rate (FDR) correction, as implemented by the Reactome online platform.

### Quantitative Reverse-Transcription Polymerase Chain Reaction

After stimulation described above, cells were harvested, and total RNA was extracted using the innuPREP RNA mini-Kit (IST) according to the manufacturer’s instructions. RNA concentration and purity were assessed using a DS-11 spectrophotometer (DeNovix). Reverse transcription into cDNA was performed using SuperScript III, RNase Out and dNTPs(all from Thermo Fisher) following the manufacturer’s protocol. For host gene expression analysis, oligo(dT) primers (Thermo Fisher, #18418020) were used for reverse transcription, whereas archaeal RNA was reverse transcribed using random hexamers (Thermo Fisher, #N8080127). Quantitative PCR was conducted using SYBR Green Master Mix on a LightCycler 480 II (both from Roche). All primers were obtained from Thermo Fisher, with their sequences provided in Supplementary Table 1 of the Supplementary Material. The cycling conditions were as follows: 95°C for 10 minutes, followed by 45 cycles of 95°C for 10 seconds, 63 - 58°C for 10 seconds with a -0.5°C/cycle gradient, and 72°C for 6 seconds. Data analysis was performed using LightCycler 480 software (v. 1.5.1), with target gene expression normalized to the reference gene HPRT. Results were expressed as n-fold induction relative to the control group.

### Flow Cytometric Analysis

For uptake experiments, PBMCs were stimulated with *M. stadtmanae* or *M. smithii* at 1 × 10^7^ cells/mL for 4 h at 37°C and 5% CO2. Unstimulated PBMCs served as controls. After stimulation, cells were stained with CD14-PE (BioLegend, #301805, 1:100) to identify monocytes. F_420_ autofluorescence of methanogenic archaea was detected using a 405 nm violet laser and a 450/50 band-pass filter, as previously described^70^. Gates were set on CD14^+^ and CD14^−^ cell populations to quantify F_420_^+^ events, representing archaea-associated monocytes. Flow cytometric acquisition was performed on a BD FACSymphony™ A3 analyzer. Imaging flow cytometry was carried out on a BD FACSDiscover™ S8 system to confirm intracellular localization of F420^+^ archaea. Propidium iodide was used for viability assessment and detected using a 561 nm yellow-green laser and a 610/20 band-pass filter. Data were analyzed using FlowJo™ software (v10.8.1), and image processing and export were performed using the BD CellView™ Lens plugin.

### Confocal Laser Scanning Microscopy

For phagocytosis assays, primary monocytes or moDCs (1 × 10^5^ cells/well) were seeded into 8-well μ-Slides (Ibidi) and equilibrated at 37 °C for 2 h. Monocytes were stimulated with whole cells of *M. stadtmanae* or *M. smithii* (1 × 10^7^ cells/mL) for 1, 2, or 4 h, whereas moDCs were stimulated under the same conditions for 4 h. Cells were then fixed with 2% paraformaldehyde, permeabilized with 0.1% Triton X-100, and blocked with 3% BSA in PBS. Archaeal cells were stained with affinity-purified rabbit antisera raised against *M. stadtmanae* or *M. smithii* (Davids Biotechnologie GmbH, 1:100), which had been generated and validated previously^57^, followed by Alexa Fluor 594-conjugated goat anti-rabbit IgG secondary antibody (1:100). Nuclei were counterstained with DAPI. Images were acquired on a Leica TCS SP5 confocal microscope using a 63× oil immersion objective. Quantification was performed in ImageJ (NIH) using the multi-point tool by counting archaeal ring-like structures in individual cells. For NF-κB p65 staining, primary monocytes (1 × 10^5^ cells/well) were seeded into 8-well μ-Slides (Ibidi), equilibrated at 37 °C for 2 h, and stimulated as indicated for 1, 2, or 4 h. Cells were then fixed with 4% paraformaldehyde, permeabilized with 0.1% Triton X-100, and blocked with 3% BSA in PBS. Cells were stained with rabbit anti-NF-κB p65 antibody (1:100, Thermo Fisher, #51-0500), followed by Alexa Fluor 546-conjugated goat anti-rabbit IgG secondary antibody (1:200, Invitrogen). Nuclei were counterstained with DAPI. Images were acquired on a Leica TCS SP5 confocal microscope using a 63× oil immersion objective and LAS AF software. Nuclear p65 fluorescence intensity was quantified in ImageJ (NIH) as the mean signal within DAPI-defined nuclear regions of interest.

### Generation of archaeal antibodies

Polyclonal antibodies against *M. smithii* and *M. stadtmanae* were generated by Davids Biotechnologie GmbH by immunizing rabbits with whole-cell biomass of the respective archaeal species. A 63-day immunization protocol with five injections was used, and antibody titers were assessed by ELISA on day 35. Affinity-purified antisera were used for all experiments in this study^57^.

### TLR8 knockout BLaER1 cells

A previously generated TLR8 knockout BLaER1 cell line was used in this study^51^. The clone was established by CRISPR/Cas9-mediated genome editing and validated previously, as described earlier^51^. TLR8 KO and corresponding control BLaER1 cells were transdifferentiated and used for stimulation experiments as indicated.

### Chromatin Immunoprecipitation (ChIP)-qPCR

ChIP-qPCR was performed as described in the Cold Spring Harbor Protocol^71^ with modifications from the fast ChIP method^72^. Briefly, monocytes stimulated with *M. stadtmanae* or *M. smithii* for 3 h were crosslinked with 1% paraformaldehyde (Thermo Fisher) for 10 min at 37°C. Glycine (0.125 M) was added to quench crosslinking. For STAT1/2 ChIP, an additional treatment with 10 mM dimethyl 3,3’-dithiobispropionimidate (DTBP, Thermo Fisher Scientific), followed by quenching with 2.5 ml of 2.5 M glycine for 5 min prior to formaldehyde crosslinking, was performed to stabilize protein-protein interactions. Cells were lysed in SDS lysis buffer (1% SDS, 10 mM EDTA, 50 mM Tris-HCl, pH 8.1), and chromatin was sheared into 200-500 bp fragments using a Bioruptor® Pico sonicator (30 sec ON/30 sec OFF, 15 cycles, 4°C). For immunoprecipitation, 10% of the chromatin lysate was saved as input control; the remainder was split 1:1 for independent STAT1 and STAT2 pulldowns (with an additional DTBP crosslinking step prior to formaldehyde), while a parallel chromatin aliquot from the same sample was used for a single NF-κB p65 pulldown (formaldehyde only). Each aliquot was incubated overnight at 4°C with one of the following antibodies conjugated to Protein A/G magnetic beads (MCE): 5 µg of anti-NF-κB p65 (Thermo Fisher, #51-0500), anti-STAT1 (Cell Signaling Technology, #9172S, 1:50 dilution), anti-STAT2 (Cell Signaling Technology, #4594S, 1:50 dilution), or Normal Rabbit IgG (Cell Signaling Technology, #2729) as a negative control. Beads were washed sequentially with low-salt (150 mM NaCl), high-salt (500 mM NaCl), and LiCl (0.25 M LiCl, 1% NP-40, 1% sodium deoxycholate, 1 mM EDTA, 10 mM Tris-HCl, pH 8.1) buffers. Crosslinks were reversed by heating at 65°C overnight in elution buffer (1% SDS, 0.1 M NaHCO), followed by RNase A and proteinase K treatment. DNA was purified using ChIP DNA Clean & Concentrator (ZYMO). qPCR was performed on both input (10%) and immunoprecipitated DNA using primers targeting NF-κB binding regions in selected Class I and Class II gene promoters. Primers were designed against upstream promoter regions based on ENCODE ChIP-seq data for p65/RelA (ENCSR000EBI) and STAT1 (ENCSR000EHK), with sequences provided in Supplementary Table 1. Enrichment was calculated as % input = 2^(ΔCt) × 100%, where ΔCt = Ct(input) − Ct(IP) - log_2_(input dilution factor).

### Transcription Factor Activity Inference and Motif Enrichment Analysis

To estimate transcription factor (TF) activities from gene expression data, we employed the *decoupleR* R package (version 2.15.0)^73^. Specifically, we utilized the Weighted Sum (WSUM) method, which integrates prior knowledge of TF-target interactions with gene expression statistics to infer TF activity scores. The TF-target interaction network was sourced from the DoRothEA regulon, a curated collection of human TF-target interactions (literature-curated resources, ChIP-Seq peaks, inference from gene expression and TF binding motif on promoters)^74^ annotated with confidence levels ranging from A (highest) to E (lowest). For this analysis, we selected interactions with confidence levels A, B, and C to ensure high reliability. To specifically dissect the contribution of the two gene modules identified in our dataset, we applied the Class I and Class II gene lists and used their log_2_ fold-change values under 4-h stimulation with *M. stadtmanae* or *M. smithii* as inputs. This allowed us to compare TF activity patterns across the two archaeal species within each gene class. To visualize these comparisons, we generated paired scatter plots (pair plots) of TF activity scores between *M. stadtmanae* and *M. smithii* for Class I and Class II genes. Motif enrichment analysis was performed using HOMER on an Ubuntu Linux platform, as previously described^75^. The Class II gene list was analyzed with HOMER findMotifs.pl, using TSS-centered regions spanning −2000 bp to +2000 bp. Known motif enrichment was assessed relative to sequence-matched background regions, and significance was reported as *P* values and Benjamini-adjusted *q* values.

### Western blot analysis

PBMC-derived monocytes were stimulated as indicated and lysed in RIPA buffer (Thermo Fisher, #89900) supplemented with protease and phosphatase inhibitors. After incubation on ice for 30 min, lysates were centrifuged at 14,000 × g for 30 min at 4°C. The supernatants were carefully transferred to fresh tubes for downstream protein quantification and Western blot analysis. Protein samples were separated using the Invitrogen NuPAGE system (Thermo Fisher) according to the manufacturer’s instructions and transferred to membranes for immunoblot analysis. Membranes were blocked for 1 h at room temperature using commercially available 1× blocking buffer prepared from a 10× stock solution (Carl Roth, # A151.4). Membranes were then incubated with primary antibodies diluted in blocking buffer overnight at 4 °C. The following primary antibodies were used: rabbit monoclonal anti-phospho-STAT1 (Tyr701) (Cell Signaling Technology, # 9167, 1:1000), mouse monoclonal antibody anti-STAT1 (Abcam, #ab281999, 1:1000), and mouse monoclonal antibody anti-β-actin (Cell Signaling Technology, #3700, 1:3000). After washing with TBST, membranes were incubated with LI-COR secondary antibodies for 1 h at room temperature at a dilution of 1:20,000: IRDye 800CW goat anti-rabbit IgG (#926-32211) for rabbit primary antibodies and IRDye 680RD goat anti-mouse IgG (#926-68020) for mouse primary antibodies. After additional washes with TBST, signals were detected using an Odyssey CLx infrared imaging system (LI-COR). Band intensities were quantified using Image Studio Lite software.

### Other R packages

Gene Set Enrichment Analysis (GSEA) was performed with *clusterProfiler* (BH correction)^76^ on preranked gene lists following *M. stadtmanae* stimulation at 4 h. Analyses focused on predefined Class I/II modules and GO categories (BP and MF). Genes were ranked by a signed statistic (log_2_FC × −log_10_ p; falling back to log_2_FC when p was unavailable). Parameters were set to minGSSize = 10, maxGSSize = 500, and 10,000 permutations with a fixed random seed. Statistical significance was defined a priori as FDR q < 0.05 together with nominal p < 0.01, and pathways were interpreted only when the leading edge subset comprised ≥10 genes. Heatmaps were generated with the *pheatmap*^77^ package. Additionally, paired dot plots, including volcano plots, were generated using the *ggplot2* package.

### Statistical Analysis

Statistical analyses were conducted using GraphPad Prism 10.4.1, except for RNA-seq and other analyses conducted in R, which are described in the corresponding Methods sections above. Statistical tests are specified in the corresponding figure legends. Exact P values are shown in the figures where applicable. To account for donor-to-donor variability, experiments were performed using primary cells from independent human donors. A batch was defined as an independent experimental run performed separately in time, typically on a different day. Depending on sample availability and assay design, each assay included 3 to 9 donors across 3 to 9 independent batches. The exact number of donors and batches used in each experiment is indicated in the corresponding figure legend.

## Results

### Shared and stimulus-specific immune transcriptional programs with distinct temporal dynamics in PBMCs stimulated by methanogenic archaea

Our previous studies confirmed that human gut-enriched methanogenic archaea, *M. stadtmanae* and *M. smithii*, activate the immune system through phagocytosis by human immune cells^49, 51^. Yet the mechanisms shaping their immunomodulatory effects remain incompletely defined. Under dose- and time-matched conditions, *M. stadtmanae* elicited significantly higher cytokine release (e.g., IL-1β, TNF-α) than *M. smithii*, indicating stimulus-specific immunostimulatory properties despite shared methanogenic metabolism^15, 49, 50^. To dissect these responses, we performed RNA-seq on human primary PBMCs from nine donors in a paired design. For each donor, cells were assigned to *M. stadtmanae*-stimulated, *M. smithii*-stimulated, or unstimulated control groups and continuously stimulated for 4 h or 24 h. This design enabled joint analysis of species effects and the temporal evolution of early (4 h) versus sustained (24 h) responses.

Because both are methanogenic archaea with shared core features, we first sought a compact overview of their common immunostimulatory footprint. A joint statistical model integrating all four stimulated conditions (4 h and 24 h, *M. stadtmanae* and *M. smithii*) identified 10,261 genes with a significant effect across the overall experimental design (adjusted p < 0.01). KEGG enrichment of these genes yielded the top 20 pathways, all with positive signed pi-scores, consistent with predominant pathway-level upregulation (Fig. 1a). These included “infection” pathways named for bacteria (e.g., Salmonella, Yersinia, Shigella, pathogenic Escherichia coli) and viruses (Epstein-Barr virus, cytomegalovirus, Kaposi sarcoma-associated herpesvirus). In KEGG, these terms chiefly denote host-side signaling and effector modules annotated in the context of infection; their enrichment in our host transcriptomes indicates that archaeal stimulation mobilizes canonical antimicrobial defense programs rather than implying the presence of these pathogens (Fig. 1a). In parallel, key innate immune signaling cascades such as Toll-like receptor, NOD-like receptor, and C-type lectin receptor pathways were significantly enriched, consistent with phagocytic uptake and pattern recognition receptor activation. Interestingly, several pathways linked to metaflammation and chronic immune modulation, including non-alcoholic fatty liver disease, lipid metabolism, and AGE-RAGE signaling, were also enriched, suggesting potential crosstalk between microbial sensing and metabolic regulation (Fig. 1a).

**Fig. 1.**
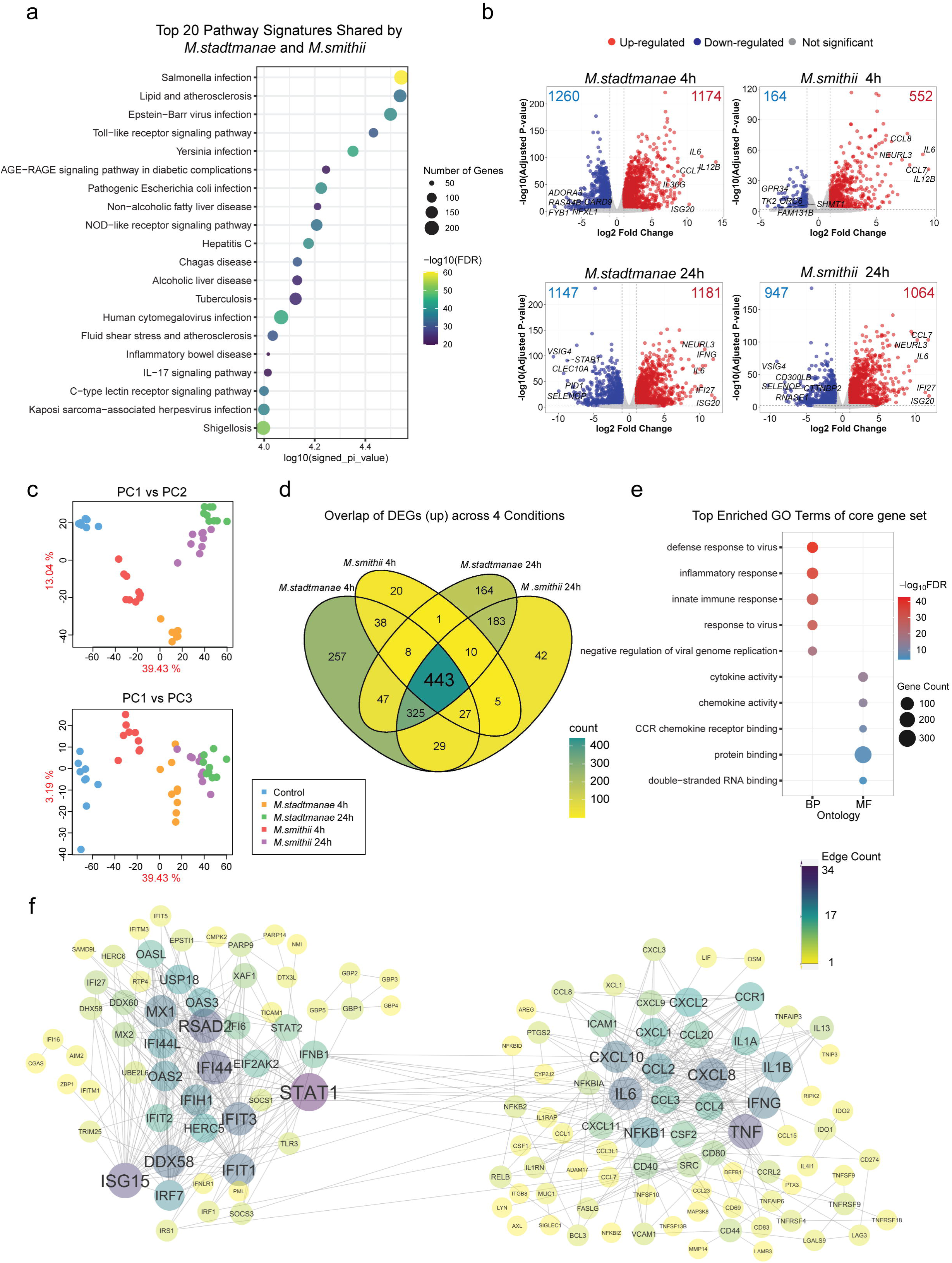
Shared and stimulus-specific immune transcriptional programs with distinct temporal dynamics in PBMCs stimulated by methanogenic archaea. **a** A bubble plot displays the top 20 KEGG pathways significantly enriched (FDR < 0.01) among genes differentially expressed in PBMCs after stimulation with *M. stadtmanae* and *M. smithii* at 4 h and 24 h; differential expression was assessed in a joint model integrating all conditions. The x-axis shows the signed pi-score (−log_10_[FDR] × sign(mean log_2_FC)), which encodes directionality (positive = net upregulation) and enrichment strength (larger magnitude = stronger significance). Bubble size indicates the number of genes mapped to each pathway, and color represents the FDR. **b** Volcano plots generated using transcriptomic data analyzed by Sleuth, with log_2_FC on the x-axis and -log_10_ p-value on the y-axis. The plots show differential gene expression in PBMCs after 4 and 24 h stimulation with *M. stadtmanae* or *M. smithii*, compared to the unstimulated control condition. Red dots indicate upregulated DEGs, blue dots indicate downregulated DEGs, and gray dots represent non-significant changes. The top 5 most upregulated and downregulated genes are labeled. The total numbers of up- and downregulated DEGs are annotated in red and blue, respectively, on the corresponding sides of each plot. **c** Principal component analysis is shown as two 2D projections of the first three components, with panels displaying PC1 versus PC2 and PC1 versus PC3. Samples are color-coded by stimulation condition, including Control, *M. stadtmanae* (4 h and 24 h), and *M. smithii* (4 h and 24 h). **d** A Venn diagram illustrates the overlap of upregulated DEGs in response to *M. smithii* and *M. stadtmanae* at 4 h and 24 h. **e** A bubble plot displays the top five enriched GO terms in the categories of BP and MF, based on the 443 core genes in response to both *M. smithii* and *M. stadtmanae* at 4 h and 24 h. BP denotes Gene Ontology Biological Process and MF denotes Gene Ontology Molecular Function. Bubble size indicates the number of core genes annotated to each term, while colour represents the FDR for enrichment significance. **f** High-confidence interactions (STRING score ≥ 0.9) among the core genes are shown, with two major clusters comprising a total of 403 nodes. Node size is proportional to the number of connected edges (degree), and node color also reflects the edge counts. Transcriptomic analyses in this figure were performed using PBMCs from 9 independent donors, with each donor representing one independent experimental batch.

We next profiled species- and time-specific transcriptional changes by differential expression analysis (adjusted P < 0.01; |log_2_fold change (FC)| ≥ 1). At 4 h, the analysis revealed a more robust response to *M. stadtmanae*, with a greater number of differentially expressed genes (DEGs) than *M. smithii*, as visualized in the volcano plots (Fig. 1b). By 24 h, DEG numbers converged: *M. stadtmanae* (1181 up, 1147 down) and *M. smithii* (1064 up, 947 down). Among the top absolute log_2_FC genes across conditions were pro-inflammatory mediators (e.g., *IL6*, *IL12B*), highlighting their consistent involvement in archaea-driven immunity (Fig. 1b). Complete results for all analyzed genes across all conditions, including log_2_FC, adjusted P-values, and joint-model P-values, are provided in Supplementary Data 1. Two-dimensional PCA of PC1 versus PC2 and PC1 versus PC3 showed a clear separation of stimulated samples from controls, especially at 24 h (Fig. 1c). Samples from the same condition clustered closely across donors. At 4 h, *M. stadtmanae* and *M. smithii* were more clearly separated from each other, while by 24 h their transcriptomic profiles were more similar, although still not fully overlapping. To assess transcriptional similarity, we generated heatmaps based on the top 100 most variable genes. The expression heatmap (Supplementary Fig. 1a, left) showed clear clustering of samples by condition, with each group forming compact clusters. Notably, *M. stadtmanae* and *M. smithii* samples at 24 h displayed highly similar expression patterns, whereas their 4 h counterparts segregated into distinct clusters with greater differences in expression amplitudes. This was further supported by the correlation heatmap (Supplementary Fig. 1a, right), which revealed strong within-group similarity, particularly among 24 h-stimulated and control samples. By contrast, pairwise cross-species correlation coefficients at 4 h remained lower, indicating more divergent early-phase responses.

Overlap analysis of upregulated DEGs across time points and species delineated shared and stimulus-specific modules (Fig. 1d). At 4 h, most *M. smithii*-induced genes (93.5%) overlapped with *M. stadtmanae* targets, though *M. stadtmanae* triggered many more DEGs overall. Notably, a conserved “core” set of 443 genes was induced under all stimulated conditions. 325 genes uniquely upregulated by *M. stadtmanae* at 4 h were also induced by *M. smithii* at 24 h, indicating a delayed response in *M. smithii*-stimulated cells. In contrast, 257 genes were exclusive to *M. stadtmanae* at 4 h, representing an early, species-dependent program. By 24 h, the overlap between the two species was substantial (961 genes), including 183 genes co-upregulated only at this later time point (Fig. 1d). Overall, these patterns indicate that the two methanogens engage a conserved core immune program with stimulus-specific kinetics and breadth. *M. stadtmanae* mobilizes a broad early module at 4 h, whereas *M. smithii* follows a delayed trajectory in which many of the same genes become induced only by 24 h. GO analysis of the 443 core genes showed strong enrichment of innate immune and inflammatory programs, with viral defense and innate immune regulation dominating Biological Process (BP) terms and cytokine/chemokine activity and receptor binding leading Molecular Function (MF) terms, indicating that archaeal stimulation engages canonical antiviral pathways (Fig. 1e). Complementary STRING protein-protein interaction analysis of the 443 core genes identified two major clusters comprising a total of 403 genes. One cluster was enriched for interferon/antiviral effectors, including *STAT1*, *STAT2*, *ISG15*, *RSAD2*, and the *IFIT* and *OAS* families, while the other contained pro-inflammatory cytokines and chemokines such as *TNF*, *IL6*, *CXCL8*, *CXCL10*, *CCL2*, and *CCL3*. These clusters were connected by a small set of bridging hubs, most notably *STAT1*, suggesting potential points of integration between antiviral and inflammatory signaling pathways (Fig. 1f). Species-specific temporal enrichments further refined these modules. The 257 *M. stadtmanae*-specific early responders were enriched for signal transduction, RNA polymerase II-mediated transcription, and histone-modifying kinase activities, consistent with rapid, chromatin-linked priming (Supplementary Fig. 1b, left). The 325 delayed responders, induced early by *M. stadtmanae* but only at 24 h by *M. smithii*, were enriched for immune and inflammatory responses, defense against viruses, apoptotic processes, and tRNA binding, consistent with a slower but comprehensive late-phase immune program (Supplementary Fig. 1b, middle). Among the 183 genes upregulated only at 24 h, metal-ion-related processes were uniquely enriched (Supplementary Fig. 1b, right). Copper and Zinc are well-established players in innate immune responses and host defense against infection^78^, suggesting that prolonged archaeal stimulation mobilizes metal - ion-dependent defense mechanisms in addition to core inflammatory and antiviral programs.

Downregulation patterns paralleled these findings. At 4 h, nearly all *M. smithii*-suppressed genes were contained within the *M. stadtmanae* set (Supplementary Fig. 2a). We observed 587 genes suppressed only at *M. stadtmanae* 4 h (not maintained at 24 h), whereas 312 genes were downregulated specifically at 24 h in both *M. stadtmanae* and *M. smithii* (Supplementary Fig. 2a), indicating a delayed suppression module. GO analysis was feasible for two downregulated subsets: *M. stadtmanae* 4 h-specific (n = 587) and 24 h-only (n = 312). The *M. stadtmanae* 4 h-specific set enriched intracellular signaling, adaptive immunity, and autophagy with lysosome/TCR complex and cytosol-nucleoplasm terms, consistent with early attenuation of signaling/immune compartments (Supplementary Fig. 2b). The 24 h-only set enriched translation and receptor-mediated endocytosis with cytosolic ribosome and extracellular exosome/space, indicating a delayed repression of protein-synthesis and secretory/extracellular pathways (Supplementary Fig. 2b). Since several monocyte subset-defining markers were among the downregulated DEGs, we examined their per-donor expression for classical (*CD14, LYZ, S100A8, S100A9*) and nonclassical (*FCGR3A/CD16, CX3CR1*) subsets. Similar reductions in these markers have been described after immune activation, reflecting transcriptional repression and receptor internalization or shedding as part of monocyte phenotypic remodeling^79^. Across donors, both archaea reduced classical and nonclassical markers, at 4 h the suppression was stronger and more consistent with *M. stadtmanae*, whereas *M. smithii* showed modest decreases that became pronounced by 24 h, at which point both stimuli converged to similar low-expression levels, with the exception of *FCGR3A/CD16*, which exhibited partial recovery at 24 h (Supplementary Fig. 2c). This indicates that archaeal stimulation is not subset-specific but instead promotes a shared reprogramming of monocytes, reducing subset-identity signatures rapidly under *M. stadtmanae* and more gradually under *M. smithii* (Supplementary Fig. 2c).

In summary, paired 4 h/24 h profiling of *M. stadtmanae* and *M. smithii* reveals a large conserved immune program, captured by a 443-gene core enriched for interferon/antiviral and inflammatory pathways, overlaid with stimulus-specific kinetics. *M. stadtmanae* elicits a more robust and earlier response at 4 h, whereas *M. smithii* follows a delayed trajectory that largely converges by 24 h. Despite this convergence, condition-specific modules persist (an early *M. stadtmanae*-biased set and genes induced by *M. smithii* only at 24 h). Together, these findings provide the first systematic, time-resolved framework for understanding how human PBMCs respond to the two dominant gut methanogens, revealing both a shared core response and species-dependent dynamics, and providing the basis for the mechanistic analyses that follow.

### Early divergence of Class I/II modules in a conserved 443-gene core across species

Previously, cytokine release assays suggested that *M. stadtmanae* is more immunogenic than *M. smithii*^49^, but whether this extends beyond a few sentinel cytokines remained unclear. To obtain a robust and comparable baseline for deeper analysis, we first defined a “conserved core” of 443 genes that were significantly upregulated under all four stimulation conditions. This core highlights transcripts repeatedly induced across donors and time points, minimizing condition-specific noise. GO enrichment of the 443-gene core was dominated by antiviral and immune pathways, which motivated a protein-level readout. Guided by these GO results, we employed a 20-plex U-PLEX™ immunoassay deliberately enriched for analytes annotated in KEGG “viral infection” pathways (e.g., CXCL10/IP-10, CCL2/MCP-1, IL-1β, IL-6, TNF-α). Supernatants from five donors were assayed at 4 h (early secretion) and 24 h (accumulated output), and 16 cytokines with robust dynamic range were carried forward. At 4 h, *M. stadtmanae* elicited markedly stronger cytokine secretion than *M. smithii* for most targets, indicating faster immune activation (Fig. 2a). This trend largely persisted at 24 h, with cytokines such as TNF-α, IL-6, and IL-12B remaining significantly higher in the *M. stadtmanae* group (Fig. 2a). However, IL-8 and CXCL10 reached similar levels under both stimuli, and notably, CCL2 was significantly higher following *M. smithii* stimulation (Fig. 2a). These findings suggest that *M. stadtmanae* does not uniformly amplify all cytokines, and in some cases, *M. smithii* may even induce stronger responses, this raised our interest in why different cytokines follow distinct patterns of regulation.

**Fig. 2.**
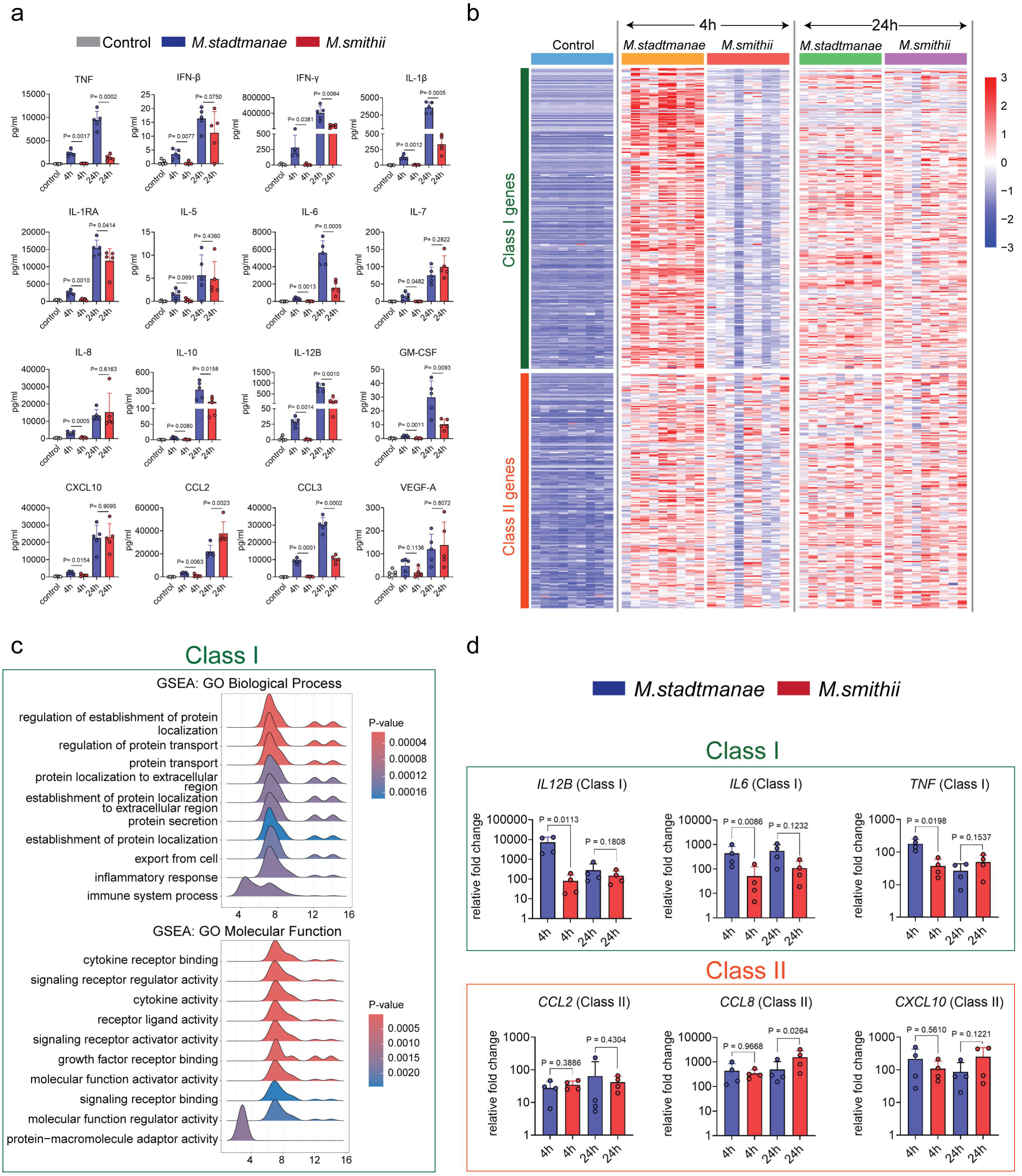
Early divergence of Class I/II modules in a conserved 443-gene core across species. **a** PBMCs were stimulated with *M. stadtmanae* or *M. smithii* for 4 h or 24 h. Cytokines and chemokines in culture supernatants were quantified using a multiplex immunoassay. Each data point represents one donor, with each donor corresponding to one independent experimental batch (n = 5). Results are shown as mean ± SD. Statistical significance between paired conditions was assessed using a two-tailed paired t-test. Exact P values are shown in the figure. **b** Heatmap displaying the normalized expression levels of Class I and Class II genes across nine individual donors (n = 9), based on transcriptome sequencing data. Samples include unstimulated controls and those stimulated with *M. stadtmanae* or *M. smithii* for 4 and 24 h. Each column corresponds to one donor under one stimulation condition. **c** GSEA was performed on Class I genes to identify enriched GO categories. The top 10 BP terms (upper panel) and MF terms (lower panel) are shown as ridge plots. Each ridge represents one GO term and shows the distribution of core enriched genes across the ranked gene list. The x-axis indicates the gene-level ranking score, and the ridge shape reflects the relative distribution of genes contributing to enrichment within each term. GO terms are displayed according to their normalized enrichment score (NES), whereas fill color denotes the enrichment p-value. **d** RT-qPCR validation of selected Class I and Class II genes identified from transcriptomic analysis. PBMCs were stimulated with *M. stadtmanae* or *M. smithii* for 4 and 24 h. Gene expression levels are presented as fold induction relative to control, normalized to the housekeeping gene *HPRT.* Each data point represents one donor and one independent experimental batch (n = 4). Data are shown as mean ± SD. Statistical significance between paired conditions was assessed using a two-tailed paired t-test. Exact *P* values are shown in the figure.

As RNA-Seq enables direct comparison of expression fold changes, we revisited the 443-gene core. Prompted by these protein-level patterns, we asked whether expression amplitudes diverge by stimulus over time. Comparing log_2_FC between species (Δ log_2_FC = log_2_FC_(*M. stadtmanae*) − log_2_FC_(*M. smithii*)), we observed the clearest separation at 4 h, where responses resolved into two modules: 247 genes more strongly induced by *M. stadtmanae* (Δ log_2_FC ≥ 1) and 195 genes similarly induced by both archaea (-1 < Δ log_2_FC < 1); only one gene favored *M. smithii* (Δ log_2_FC ≤ -1). By 24 h, 401/443 genes fell within -1 < Δ log_2_FC < 1, indicating substantial convergence. Because this early-phase divergence was most apparent at 4 h, and largely obscured by later convergence, we centered subsequent analyses on this time point and formally defined two modules: Class I (247 genes), more strongly induced by *M. stadtmanae* (Δ log_2_FC ≥ 1), and Class II (195 genes), similarly induced by both archaea (-1 < Δ log_2_FC < 1). For clarity, “Class I/II” are operational labels for these expression-defined gene modules and are unrelated to MHC class I/II antigen presentation. Detailed lists of these gene classes and Δlog_2_FC are provided in Supplementary Data 2. Heatmaps of Class I/II gene expression across nine donors (across sample normalized expression) showed the clear 4 h pattern (Fig. 2b): Class I genes were more strongly induced by *M. stadtmanae* than by *M. smithii*, whereas Class II genes were expressed at comparable levels under both stimuli. By 24 h, expression of both classes largely converged. To probe the functional implications of this two-module classification, we next examined the pathway usage of Class I and Class II. Gene set enrichment analysis (GSEA) on ranked lists showed Class I enrichment for protein transport/extracellular localization and cytokine activity/receptor binding (Fig. 2c), whereas Class II lacked significant enrichment (P > 0.01), suggesting more dispersed functions. To validate RNA-Seq findings, we selected representative genes for qPCR: *IL12B*, *IL6*, and *TNF* for Class I, and *CCL2*, *CCL8*, and *CXCL10* for Class II. *IL12B* and *IL6* were chosen as they represented the top two most strongly induced Class I genes at 4 h for both archaea, while *TNF*, although not among the top-ranked by fold change, was included due to its central role in monocyte activation and inflammatory signaling. For Class II, *CCL8* was selected as the top-ranked induced gene in both conditions, while *CCL2* and *CXCL10* were among the top 10 regulated genes and, together with *CCL8*, represent the top three chemokines at this time point. At 4 h, qPCR confirmed that Class I genes were significantly more upregulated by *M. stadtmanae*, while Class II genes showed no significant difference between stimuli (Fig. 2d). At 24 h, expression levels of most genes equalized, although *CCL8* remained higher in the *M. smithii* condition (Fig. 2d). Together, these data show that *M. stadtmanae* preferentially upregulates a defined early subset (Class I), rather than driving a globally stronger module, while a large Class II subset is stimulus independent. This Class I/II framework, introduced here from the 443-gene core, provides the organizing principle for the mechanistic analyses that follow.

### *M. stadtmanae* is more efficiently phagocytosed than *M. smithii*, with stronger Class I responses in monocytes and moDCs

Monocytes are the primary phagocytic cells within the immune system, exhibiting greater phagocytic capacity than other peripheral immune cell types^80^. Our previous work showed that immune activation by *M. stadtmanae* and *M. smithii* requires phagocytic uptake and intracellular processing^49^. Given that *M. stadtmanae* induces a more rapid and robust transcriptional response than *M. smithii* under short-term (4 h) stimulation, we hypothesized that it may be taken up more efficiently by immune cells, enabling earlier and stronger recognition and signaling. Before testing this directly, we first excluded the alternative possibility that instability of the archaea in culture results in the extracellular release of immunostimulatory material independently of cellular uptake. No archaeal signal was detected in rigorously cleared cell-free supernatants by species-specific RT-qPCR (Supplementary Fig. 3a), and these supernatants failed to induce measurable cytokine release in responder cells (Supplementary Fig. 3b), indicating that under our experimental conditions the archaea remained largely intact and did not release appreciable immunostimulatory material into the extracellular medium.

Given their distinct transcriptional effects, we next asked whether *M. stadtmanae* and *M. smithii* differ in cellular uptake by PBMCs. We first used coenzyme F_420_ autofluorescence as a rapid label-free screening readout. F_420_ is a fluorescent cofactor conserved in methanogenic archaea and essential for methanogenesis^81^, and has previously been used to isolate *M. smithii* from human samples^82, 83^. Before quantifying cellular uptake, we first determined the basal F_420_ fluorescence of free archaeal cells under identical acquisition settings and found that free *M. smithii* displayed significantly higher basal F_420_ fluorescence than free *M. stadtmanae* (Supplementary Fig. 4a, b). We next focused on CD14^+^ monocytes and, after 4 h stimulation, quantified both the proportion of F_420_-positive cells and the archaeal F_420_ intensity on a per-cell basis. CD14^+^ monocytes showed both a significantly higher fraction of F_420_-positive cells (Supplementary Fig. 5a, c) and a higher normalized per-cell F_420_ signal in the *M. stadtmanae* condition than in the *M. smithii* condition (Supplementary Fig. 5a, d). By contrast, CD14^−^ cells showed no corresponding increase in archaeal association (Supplementary Fig. 5b, e). Consistent with these findings, imaging flow cytometry revealed stronger and more abundant intracellular F_420_ puncta in *M. stadtmanae*-stimulated cells at 4 h (Supplementary Fig. 5f).

We therefore next sought to validate this uptake difference directly in monocytes and to capture its early dynamics using an independent antibodies-based approach. To this end, we performed immunofluorescence microscopy with previously validated species-specific antibodies against *M. stadtmanae* and *M. smithii*^57^. Primary CD14^+^ monocytes were isolated from PBMCs and then stimulated with whole cells of either archaeon for 1, 2, or 4 h to capture early uptake kinetics. Distinct red fluorescent ring-like archaeal structures were detected within stimulated cells, consistent with internalized archaea (Fig. 3a; Supplementary Fig. 6). No such structures were observed in unstimulated control cells. Quantification confirmed that *M. stadtmanae* was taken up both earlier and more efficiently than *M. smithii* (Fig. 3b). Already at 1 h, the uptake frequency, measured as the percentage of cells containing at least one internalized archaeal cell, was markedly higher for *M. stadtmanae* than for *M. smithii* (47.66% vs 7.32%). The same pattern was observed at 2 h (78.85% vs 13.64%) and 4 h (83.08% vs 31.04%). In addition, the uptake load, quantified as the number of internalized archaea per phagocytic cell, was consistently higher for *M. stadtmanae* than for *M. smithii* at all time points examined (Fig. 3b). Together, these independent imaging analyses validated CD14^+^ monocytes as the principal uptake population and showed that *M. stadtmanae* is internalized earlier and more efficiently than *M. smithii*. We next asked whether the transcriptional patterns observed in PBMCs could be recapitulated in purified CD14^+^ monocytes. RT-qPCR on representative genes (Class I: *IL12B*, *IL6*, *TNF*; Class II: *CCL2*, *CCL8*, *CXCL10*) and ELISA on an overlapping but non-identical panel (Class I: IL-6, TNF-α, IL-1β; Class II: CCL2, CXCL10) showed that Class I genes were more strongly induced by *M. stadtmanae*, whereas Class II genes were comparably induced by both species (Fig. 3c, d), consistent with the PBMCs patterns. To test whether this stronger Class I response depends on more efficient uptake, we inhibited phagocytosis with cytochalasin D and found that IL-6 and IL-1β responses to both archaea were strongly suppressed. Importantly, the difference between *M. stadtmanae* and *M. smithii* seen under uninhibited conditions was no longer evident after inhibition (Supplementary Fig. 5g). This indicates that the stronger Class I response to *M. stadtmanae* is largely driven by more efficient uptake.

**Fig. 3.**
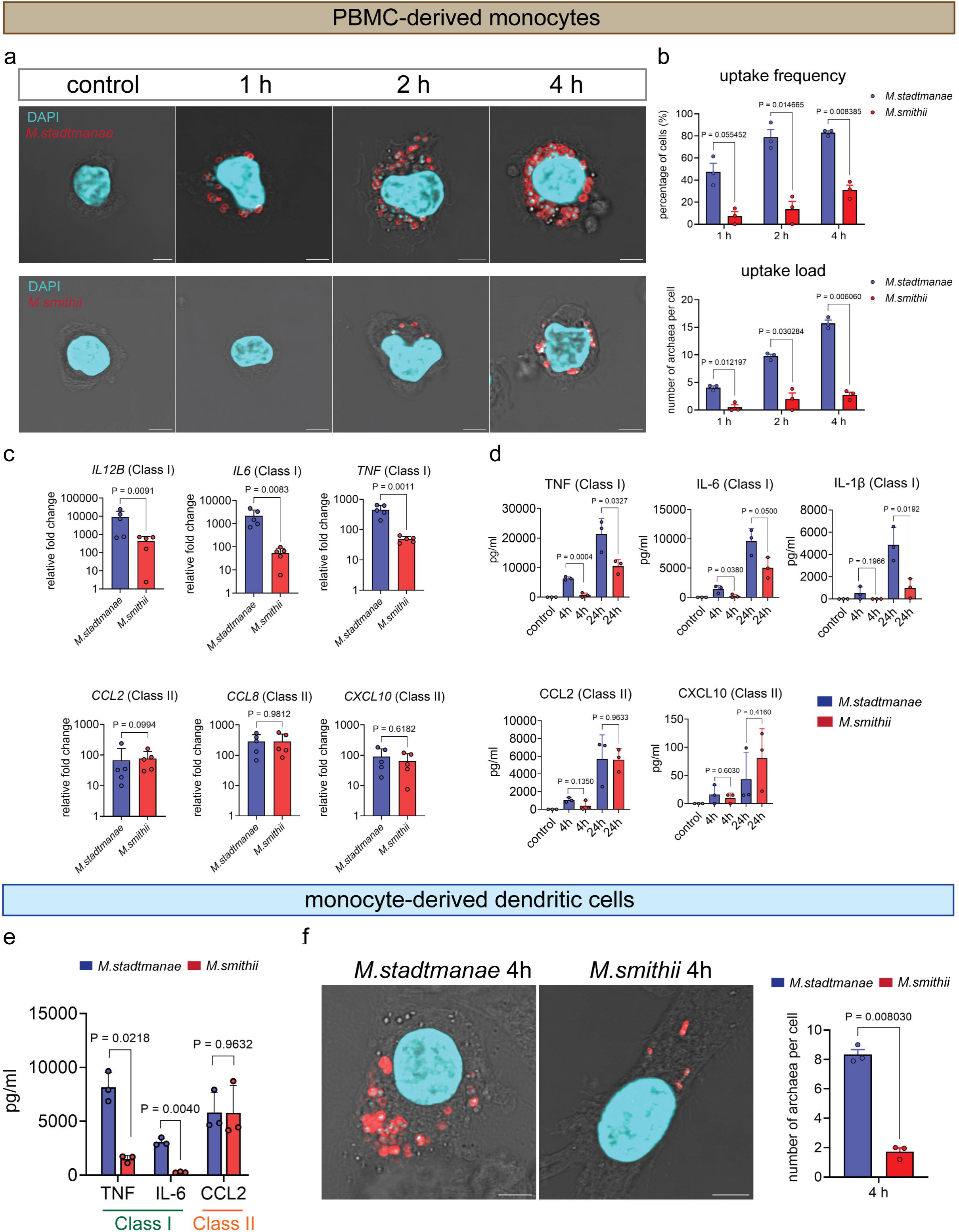
*M. stadtmanae* is more efficiently phagocytosed than *M. smithii* by CD14^+^ monocytes and moDCs. **a** Representative merged confocal images of primary monocytes stimulated with *M. stadtmanae* or *M. smithii* for 1, 2, or 4 h, as indicated. Archaea were visualized by species-specific immunofluorescence staining and appear as red fluorescent ring-like structures. Nuclei were counterstained with DAPI (cyan). Phase-contrast images are included to show cell morphology and outline. Images were acquired with a 63× oil immersion objective. Displayed images represent enlarged views of selected regions to better visualize archaea-associated signals. Scale bar = 5 µm. **b** Quantification of archaeal uptake in primary monocytes after stimulation with *M. stadtmanae* or *M. smithii* for 1, 2, or 4 h. Upper panel, uptake frequency, defined as the percentage of cells containing at least one internalized archaeal cell. Lower panel, uptake load, defined as the mean number of internalized archaea per cell, with each ring-like structure scored as one archaeal cell. For each condition, 30-50 cells were analyzed from 3 independent donors. For each donor, data were first summarized across the analyzed cells. In the upper panel, each dot represents the uptake frequency for one donor, and in the lower panel, each dot represents the mean uptake load for one donor and one independent experimental batch (n = 3). Data are shown as mean ± SEM. Statistical significance was assessed using a two-tailed paired test. P values are indicated in the figure. **c** Gene expression levels of representative Class I and Class II genes were validated by RT-qPCR in PBMC-derived monocytes stimulated with *M. stadtmanae* or *M. smithii* for 4 h. Expression is shown as fold induction relative to control, normalized to *HPRT*. Each data point represents one donor and one independent experimental batch (*n* = 5). Data are shown as mean ± SD. Statistical significance between paired conditions was assessed using a two-tailed paired t-test. Exact *P* values are shown in the figure. **d** ELISA was used to quantify TNF and IL-6 (encoded by representative Class I genes), CCL2 and CXCL10 (encoded by a representative Class II genes) in the supernatants of PBMC-derived monocytes stimulated with *M. stadtmanae* or *M. smithii* for 4 h or 24 h. Each data point represents one donor and one independent experimental batch (*n* = 3). The plots display the mean ± SD. Statistical significance was determined using a two-tailed paired t-test. Exact *P* values are shown in the figure. **e** ELISA was used to measure the secretion of TNF and IL-6 (encoded by Class I genes) and CCL2 (encoded by a Class II gene) in supernatants collected from moDCs stimulated with *M. stadtmanae* or *M. smithii* for 24 h. Each data point represents one donor and one independent experimental batch (*n* = 3). The plots display the mean ± SD. Statistical significance was determined using a two-tailed paired t-test. Exact *P* values are shown in the figure. **f** Left panel: representative confocal images and quantification of archaeal uptake in moDCs after stimulation with *M. stadtmanae* or *M. smithii* for 4 h. Archaea were visualized by species-specific immunofluorescence staining and appear as red fluorescent ring-like structures. Nuclei were counterstained with DAPI (cyan), and phase-contrast images show cell morphology and outline. Images were acquired with a 63× oil immersion objective. Displayed images represent enlarged views of selected regions to better visualize archaea-associated signals. Scale bar = 5 µm. Right panel, quantification of the number of internalized archaea per cell, with each red ring-like structure counted as one archaeal cell. For each condition, 30-61 cells from 3 independent donors/batches were analyzed. For each donor, data were first summarized across the analyzed cells. Each dot represents the mean uptake load for one donor and one independent experimental batch (n = 3). Data are shown as mean ± SEM. Statistical significance was assessed using a two-tailed paired test. Exact P values are shown in the figure.

Since previous studies have shown that *M. stadtmanae* induces higher cytokine release than *M. smithii* in moDCs^49^, we next investigated whether enhanced uptake by moDCs contributes to this difference. To generate moDCs, monocytes were differentiated with GM-CSF and IL-4. Based on earlier findings, we focused our ELISA analysis on TNF, which was reported to be strongly induced by *M. stadtmanae*^49^. In addition, we included IL-6 to extend the panel of Class I cytokines, and CCL2 as a representative Class II protein not previously examined. ELISA confirmed significantly higher TNF and IL-6 levels upon *M. stadtmanae* stimulation, whereas CCL2 secretion was comparable between the two species (Fig. 3e), mirroring the patterns observed in PBMCs and monocytes. Importantly, after 4 h stimulation, *M. stadtmanae* again showed markedly higher uptake load than *M. smithii*, with more internalized archaeal cells detected per moDC (Fig. 3f; Supplementary Fig. 7).

Together, these data identify CD14^+^ monocytes as the principal archaeal uptake population within PBMCs, show that *M. stadtmanae* is internalized earlier and more efficiently than *M. smithii*, and indicate that this uptake difference is associated with stronger Class I responses in monocytes and moDCs.

### Temporal profiling reveals dynamics of Class I and Class II genes induction by *M.stadtmanae and M.smithii*

Based on our earlier uptake measurements showing significantly earlier internalization of *M. stadtmanae* than *M. smithii*, we reasoned that differences during the earliest stimulation phase could contribute to the stronger Class I induction observed with *M. stadtmanae*. However, this uptake difference did not readily explain why Class II expression was already similar at 4 h. We therefore hypothesized that the two species differ mainly in the kinetics of Class II induction, with *M. stadtmanae* triggering an earlier response and *M. smithii* showing delayed accumulation that narrows the difference over time. To capture these early dynamics more precisely, we extended the time course to 1, 2, 4, and 6 h. We then quantified temporal expression of representative Class I (*IL12B, IL6, TNF*) and Class II (*CCL2, CCL8, CXCL10*) genes at these times.

For selected Class I genes, *M. stadtmanae* stimulation induced a sharp increase in expression between 1 and 2 h, followed by sustained high levels through 6 h (Fig. 4). A slight decrease was observed for *TNF* at 6 h, but expression remained elevated overall. In contrast, *M. smithii* induced a gradual increase in expression at all time points, yet levels remained substantially lower than those elicited by *M. stadtmanae* (Fig. 4). For Class II genes, the difference between the two stimuli was mainly kinetic. At 1 and 2 h, *M. stadtmanae* induced higher transcript levels than *M. smithii*, most clearly for *CCL2* and *CCL8*, which were already significantly upregulated at 1 h (Fig. 4). Expression then increased further under both conditions, but with different trajectories. Following *M. stadtmanae* stimulation, transcript levels rose earlier and then increased more gradually. By contrast, *M. smithii* induced a delayed but steeper increase, such that expression approached *M. stadtmanae* levels by 4 h and, for *CCL2*, exceeded them by 6 h (Fig. 4). Notably, all three Class II genes continued to rise between 4 and 6 h, indicating that 4 h does not represent the maximal Class II response under either condition. These results support the rationale for extending the early time course and show that Class I and Class II modules differ in temporal organization: Class I genes retain amplitude differences across time, whereas Class II genes differ primarily in onset and become more similar at later time points.

**Fig. 4.**
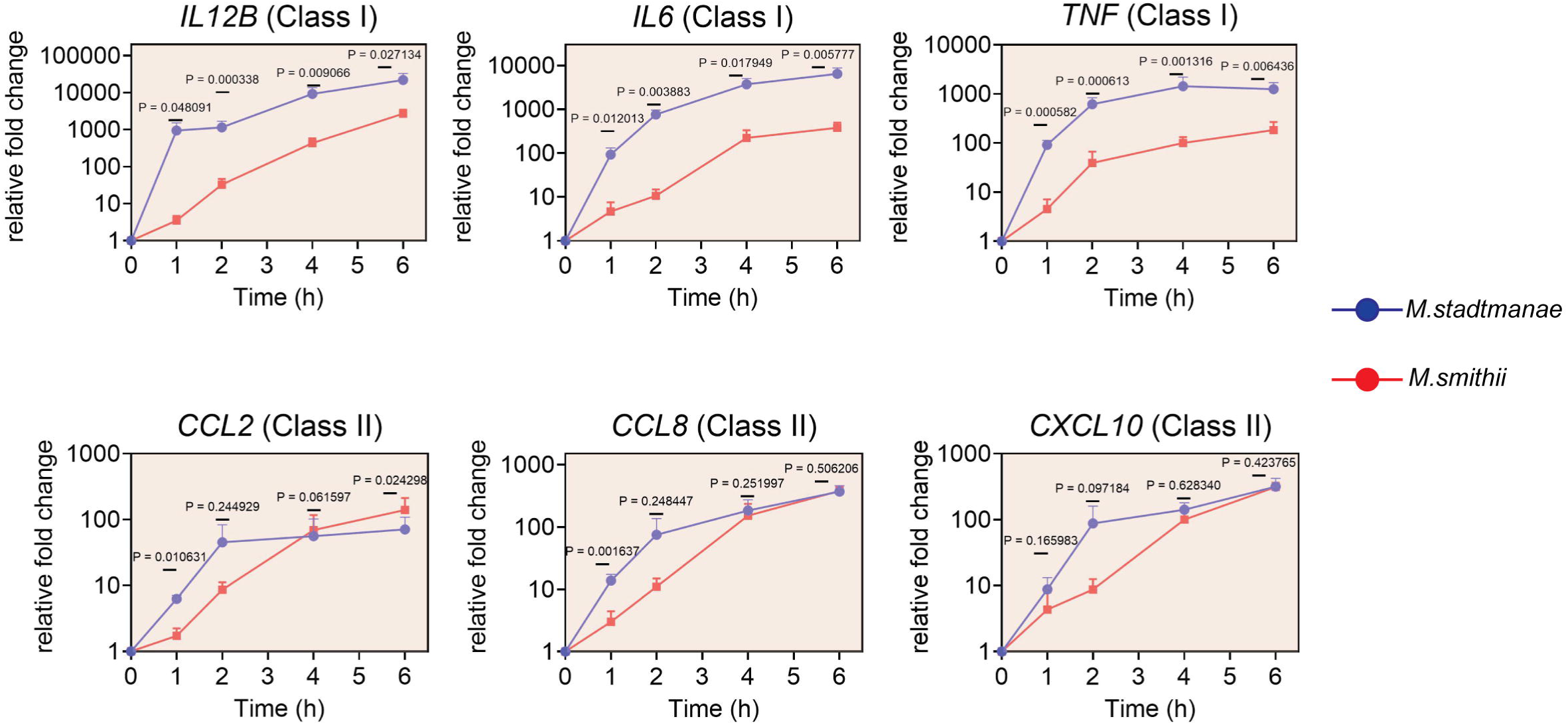
Temporal profiling reveals dynamics of Class I and Class II gene induction by *M. stadtmanae* and *M. smithii*. RT-qPCR analysis of selected Class I and II genes in PBMC-derived monocytes stimulated with *M. stadtmanae* or *M. smithii* for 1, 2, 4, and 6 h. Gene expression levels are presented as fold induction relative to control, normalized to HPRT. Data are shown as mean ± SEM. Each data point represents one donor and one independent experimental batch (n = 4). Statistical significance was assessed using multiple paired t-tests to compare *M. stadtmanae* and *M. smithii* at each time point. Exact P values are shown in the figure.

### Monocyte responses to archaea are predominantly TLR8-dependent with residual TLR8-independent signatures

Given the early kinetic divergence between Class I and Class II, we asked whether upstream receptor wiring could explain these patterns. Prior work showed that *M. stadtmanae*-induced responses in human immune cells are largely mediated through archaeal RNA sensing by endosomal TLR8^51^. However, that study was limited to a few cytokines, leaving open whether broader gene modules, including Class I and Class II genes, share this dependence or engage TLR8-independent mechanisms. To address this, we used CU-CPT9a, a selective TLR8 antagonist that locks the receptor in its resting dimeric state and blocks ligand-induced NF-κB signaling without affecting other TLRs^84–86^. As pathway controls, we stimulated cells with agonists for distinct TLRs, including LPS for TLR4, Pam3CSK4 for TLR2, and the TLR8-selective agonist TL8-506. CU-CPT9a selectively suppressed TL8-506 responses while leaving LPS- and Pam3CSK4-driven activation intact, supporting functional specificity for TLR8 under our assay conditions (Supplementary Fig. 8a). PBMC-derived monocytes were used as the principal responding cell type to minimize confounding from other leukocyte subsets. Cells were pretreated with CU-CPT9a for 1 h and then stimulated with *M. stadtmanae*, *M. smithii*, or TL8-506 for 4 h. qPCR confirmed robust inhibition of TL8-506-induced gene expression, with reduction rates ranging from 84.7% to 98.9%. Importantly, both Class I and Class II genes were strongly suppressed by the TLR8 antagonist, and stimulation with archaeon yielded similar results, with most transcript levels reduced to near baseline (Fig. 5a), suggesting that TLR8 is the dominant upstream sensor routing both Class I and Class II responses to archaeal stimulation.

**Fig. 5.**
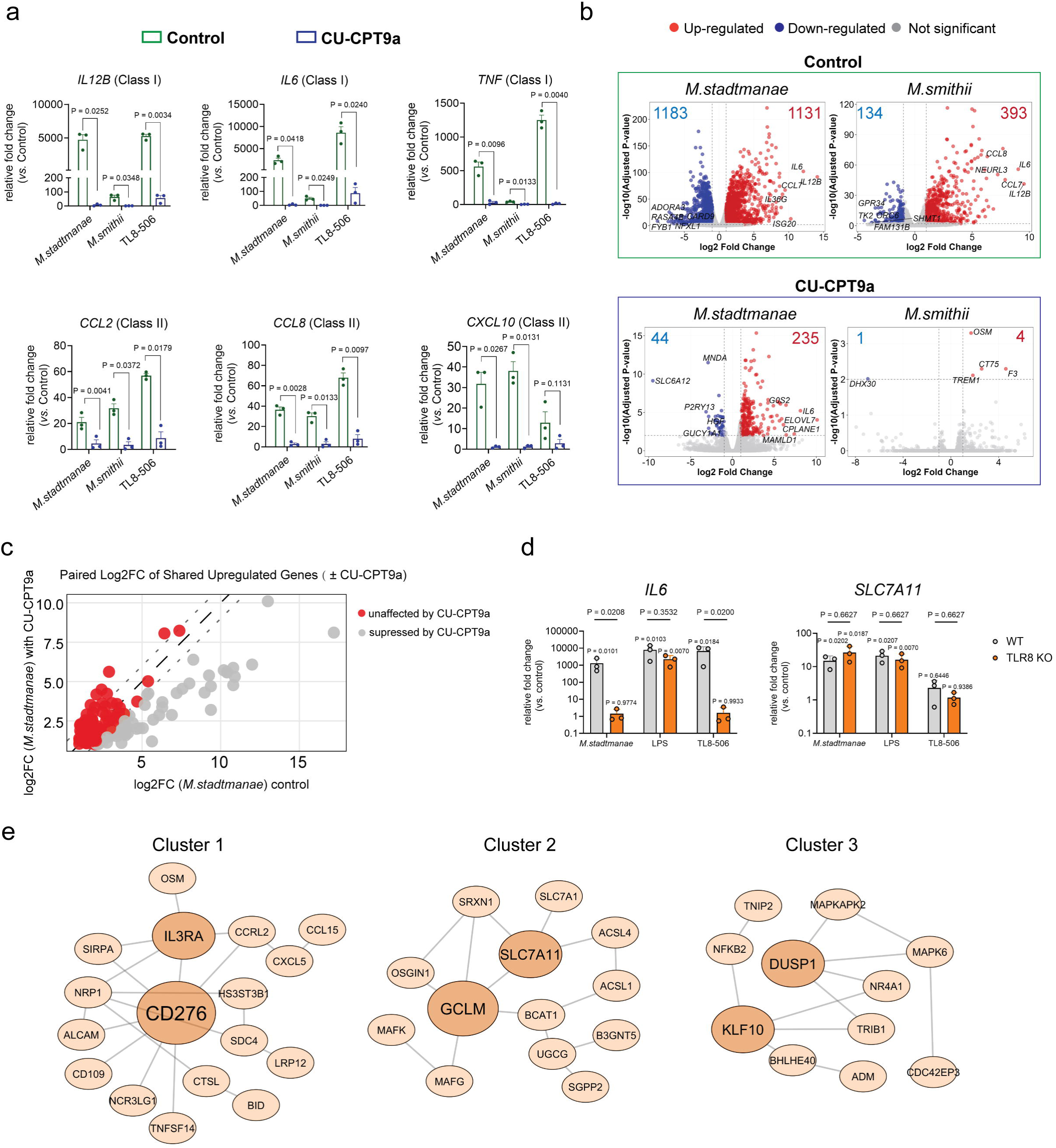
Monocyte Responses to Archaea Are Predominantly TLR8-Dependent with Residual TLR8-Independent Signatures. **a** PBMC-derived monocytes were stimulated for 4 h with *M. stadtmanae*, *M. smithii*, or the TLR8 agonist TL8-506 (1 μg/mL), with or without pre-treatment using the TLR8 inhibitor CU-CPT9a. Expression levels of representative Class I and Class II genes were measured by RT-qPCR and normalized to *HPRT*. Data are presented as fold induction relative to unstimulated controls. Each data point represents an individual donor and one independent experimental batch (*n* = 3); bars indicate mean ± SD. Statistical significance between paired conditions was assessed using a two-tailed paired *t*-test. Exact P values are shown in the figure. **b** Volcano plots were generated from RNA-seq data of PBMC-derived monocytes using Sleuth. For each condition, log_2_FC (x-axis) was plotted against -log_10_ *p*-value (y-axis). The *M. stadtmanae* and *M. smithii* groups (upper panels) were compared to the unstimulated control, while the *M. stadtmanae* +[CU-CPT9a and *M. smithii* + CU-CPT9a groups (lower panels) were compared to the CU-CPT9a-only control. Red dots represent upregulated genes, blue dots represent downregulated genes, and gray dots denote non-significant changes. The five most upregulated and five most downregulated genes (by log_2_FC) are labeled. Total numbers of DEGs for each comparison are indicated in boxed annotations. **c** Dot plot showing paired log2FC values of 171 shared upregulated genes. Each dot represents one DEG detected under both *M. stadtmanae* stimulation alone and *M. stadtmanae* stimulation in the presence of CU-CPT9a. The x-axis shows the log2FC under *M. stadtmanae* stimulation alone, and the y-axis shows the corresponding log2FC in the presence of CU-CPT9a. The long dashed diagonal line (y = x) indicates equal expression between conditions. Two additional short dashed lines, drawn parallel to the diagonal at ±1 log2FC difference, define two response categories based on CU-CPT9a sensitivity, where Δlog2FC is defined as log2FC in the absence minus log2FC in the presence of CU-CPT9a: (1) gray dots, 64 genes markedly reduced by CU-CPT9a (Δlog2FC ≥ 1); and (2) red dots, 107 genes largely unaffected by CU- CPT9a (Δlog2FC < 1). **d** RT-qPCR analysis of *IL6* and *SLC7A11* expression in WT and TLR8 KO BLaER1 cells following stimulation as indicated. TL8-506 and LPS were included to confirm selective loss of TLR8 signaling in TLR8 KO cells. Data are shown as mean ± SD from three independent experiments. Statistical significance was assessed by one-way ANOVA with Dunnett’s multiple comparisons test for comparisons of each stimulated group with its corresponding unstimulated control within the same genotype. These P values are indicated above the bars. Comparisons between WT and TLR8 KO cells under each stimulation condition were performed using multiple paired t-tests, and the corresponding P values are indicated above the connecting lines. **e** Interaction network analysis of the 107 genes that remained upregulated in the presence of CU-CPT9a. High-confidence interactions (STRING score ≥ 0.9) were retrieved from STRING and visualized in Cytoscape, revealing three connected clusters.

To determine whether this inhibitory effect extended beyond the selected genes and to obtain a global view of transcriptional regulation, we next performed RNA-seq on PBMC-derived monocytes from three donors stimulated for 4 h with each archaeon, with or without CU-CPT9a. Using |log2FC| ≥ 1 and adjusted p < 0.01, *M. stadtmanae* induced more DEGs than *M. smithii* at 4 h (Fig. 5b), consistent with PBMCs. While most *M. smithii*-upregulated genes overlapped with the *M. stadtmanae* response (Supplementary Fig. 8b), *M. stadtmanae* also uniquely induced 778 upregulated genes, mirroring the combination of shared and stimulus-specific programs observed in PBMCs (Supplementary Fig. 8b). TLR8 inhibitor CU-CPT9a alone did not alter basal transcription (Supplementary Fig. 8c). Under inhibition, the *M. smithii* signature was nearly eliminated, with only 4 DEGs remaining, whereas *M. stadtmanae* retained a largely reduced but still appreciable DEG set (Fig. 5b). Complete results for all analyzed genes across all conditions are provided in Supplementary Data 3. Hierarchical clustering and PCA provided a global view: archaeal stimulation produced profiles clearly distinct from controls, whereas TLR8 inhibition shifted samples toward controls (Supplementary Fig. 8d). Correlation analysis mirrored this pattern, that is *M. smithii* + CU-CPT9a nearly overlapped with controls (consistent with the near-complete loss of DEGs), while *M. stadtmanae* + CU-CPT9a showed only partial reversion with a residual signature (Supplementary Fig. 8e).

We next focused on the residual transcriptional response under TLR8 blockade. Although CU-CPT9a markedly reduced the *M. stadtmanae* response, 235 genes were still significantly upregulated (Fig. 5b). Of these 235 genes, 171 were significantly upregulated by *M. stadtmanae* both with and without inhibitor (Supplementary Fig. 8f). Among the 171 shared genes, two response categories emerged based on CU-CPT9a sensitivity. When defined as the difference between log2FC in the absence and presence of CU-CPT9a, Δlog2FC separated 64 genes that were markedly reduced by the inhibitor (Δlog2FC ≥ 1) but remained differentially expressed, from 107 genes that were largely unaffected (Δlog2FC < 1) (Fig. 5c).

Pharmacologic inhibition alone could not determine whether residual expression reflected incomplete blockade of TLR8 signaling or genuine TLR8-independent regulation. We therefore complemented the inhibitor-based analysis with a genetic loss-of-function approach using transdifferentiated BLaER1 monocytes. TLR8 BLaER1-knockout cells failed to respond to TL8-506 while retaining responsiveness to LPS, confirming selective loss of TLR8 signaling (Fig. 5d). We then examined representative genes from the two CU-CPT9a-defined categories under *M. stadtmanae* stimulation. *IL6*, from the inhibitor-sensitive group, was largely reduced by CU-CPT9a but remained above the DEG cutoff because of its very strong induction. Notably, *IL6* expression was abolished in TLR8 knockout cells and returned to basal levels. In contrast, *SLC7A11*, a representative gene from the inhibitor-insensitive group, was unchanged by both CU-CPT9a treatment and TLR8 knockout (Fig. 5d). These results show that residual expression after CU-CPT9a treatment can still be compatible with full TLR8 dependence, as seen for *IL6*, whereas inhibitor-insensitive genes such as *SLC7A11* are consistent with TLR8-independent regulation.

We next analyzed the 107 genes that remained upregulated in the presence of the TLR8 antagonist. Interaction network analysis revealed that a subset of these transcripts formed three connected clusters. Cluster 1 was centered on *CD276* and *IL3RA*. Cluster 2 was enriched for redox-associated genes and was organized around the *SLC7A11–GCLM* axis. In agreement with this redox signature, Reactome pathway analysis of the CU-CPT9a-insensitive genes showed strong enrichment of NFE2L2-related pathways, with the top hits including “NFE2L2 regulating antioxidant/detoxification enzymes” and the “KEAP1–NFE2L2 pathway,” pointing to a potential contribution of NFE2L2-driven transcription to these TLR8-independent responses (Supplementary Fig. 8g). Cluster 3 comprised an immediate-early signaling module featuring *DUSP1*.

Together, these data define archaeal-induced monocyte responses as predominantly TLR8-driven, with a smaller set of genes remaining TLR8-independent.

### Selective low-dose activation of Class II genes by archaeal RNA

Our previous work had established that immune responses to *M. stadtmanae* are largely mediated through RNA-dependent TLR8 sensing^51^. In the present study, our inhibitor experiments further showed that responses to both *M. stadtmanae* and *M. smithii* depend predominantly on TLR8. This raised the question of whether intrinsic differences between the RNAs of the two species contribute to the distinct balance of Class I and Class II responses they induce.

We therefore used DOTAP-mediated delivery of total archaeal RNA to compare RNA-intrinsic immunostimulatory activity under standardized delivery conditions. DOTAP is a widely used cationic lipid reagent for intracellular RNA delivery and has been used in functional TLR7/8 RNA-sensing assays to enable RNA ligand access to intracellular RNA-sensing pathways.^51, 87–90^. DOTAP at the working concentration used for RNA delivery (10 µg/ml) did not reduce cell viability (Supplementary Fig. 9a, b). Naked archaeal RNA was inactive, whereas DOTAP-complexed RNA induced robust cytokine production, indicating that efficient delivery was required in this system (Supplementary Fig. 9c). This stimulatory activity was completely lost after RNase A treatment, while DOTAP-complexed archaeal DNA, tested at the same concentration as archaeal RNA, remained inactive under the same conditions (Supplementary Fig. 9d), showing that RNA was responsible for the activity. Moreover, cytokine induction by DOTAP-complexed archaeal RNA was completely abolished by the selective TLR8 antagonist CU-CPT9a (Supplementary Fig. 9c). Together, these findings show that the response operates through an RNA-TLR8 axis.

Under equal-input, equal-delivery conditions, RNA from *M. stadtmanae* and *M. smithii* induced comparable expression of both Class I and Class II genes at the transcript level, as measured by RT-qPCR (Fig. 6a), and similarly comparable levels of IL-6, IL-1β, TNF, and CCL2 at the protein level, as measured by ELISA (Fig. 6b). These findings indicate that RNA from *M. stadtmanae* and *M. smithii* has comparable intrinsic stimulatory activity under equal-input, equal-delivery conditions. Prior whole-cell time-course data add a second piece: despite lower early uptake, *M. smithii* still achieves Class II expression comparable to *M. stadtmanae*.

**Fig. 6.**
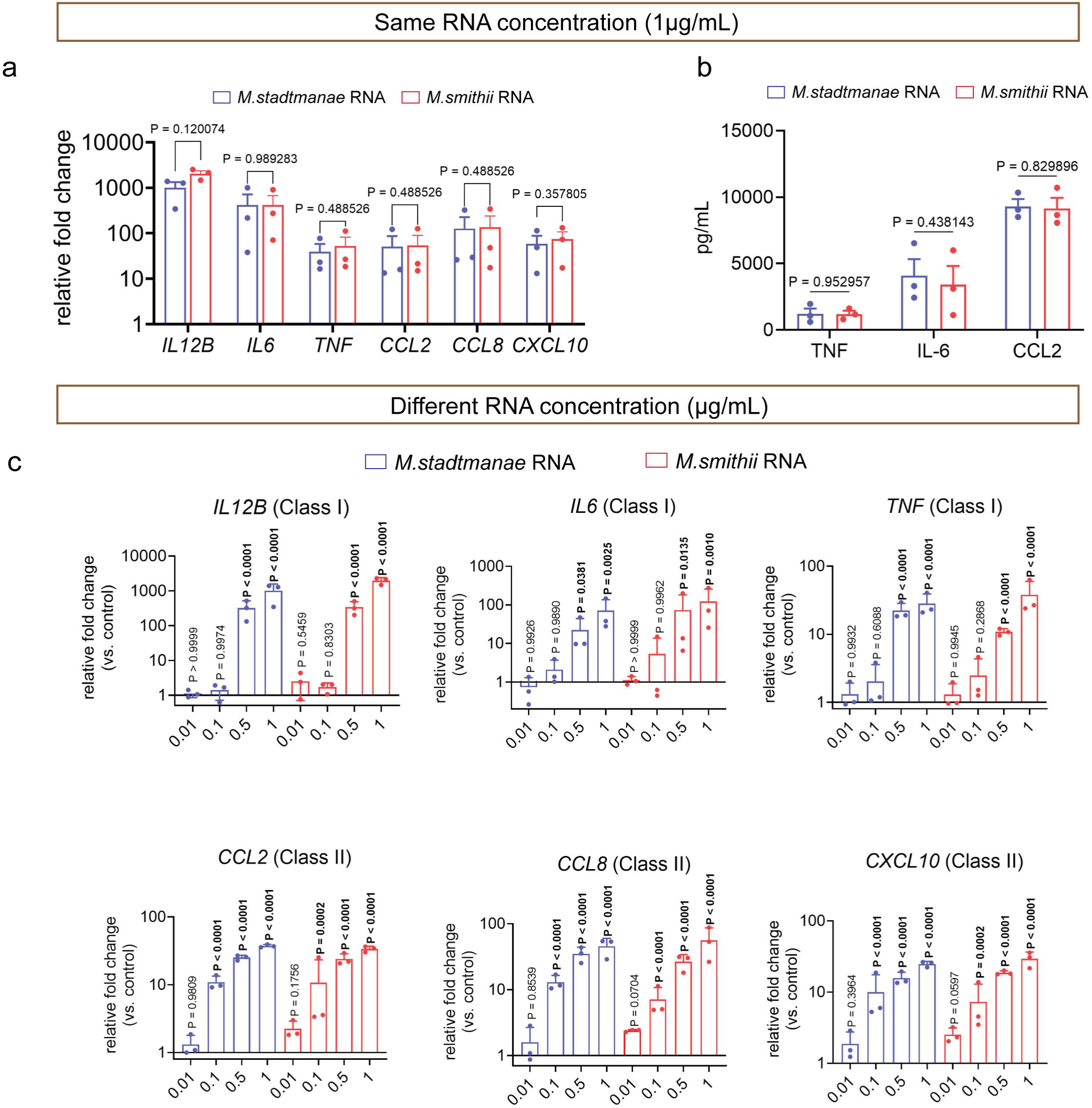
Selective Low-Dose Activation of Class II Genes by Archaeal RNA. **a** PBMC-derived monocytes were stimulated for 4 h with total RNA (1 μg/mL) extracted from *M. stadtmanae* or *M. smithii*, followed by RT-qPCR quantification of selected Class I and Class II genes. Gene expression levels are presented as fold induction relative to control, normalized to *HPRT*. Each dot represents one donor, corresponding to one independent experimental batch (n = 3). Bars indicate mean ± SEM. Statistical significance was assessed using a two-tailed paired *t*-test. Exact P values are indicated above the connecting lines. **b** ELISA analysis of TNF, IL-6, and CCL2 in supernatants from PBMC-derived monocytes stimulated for 4 h with total RNA (1 μg/mL) extracted from *M. stadtmanae* or *M. smithii*. Each dot represents one donor, corresponding to one independent experimental batch (n = 3). Bars indicate mean ± SEM. Statistical significance was assessed using multiple paired t-tests. Exact P values are indicated above the connecting lines. **c** RT-qPCR analysis of selected Class I and Class II gene expression in PBMC-derived monocytes stimulated for 4 h with varying concentrations (0.01, 0.1, 0.5, and 1 μg/mL) of total RNA extracted from *M. stadtmanae* or *M. smithii*. Gene expression levels are presented as fold induction relative to control, normalized to *HPRT*, with the y-axis shown on a log_10_ scale to reflect the magnitude of induction. Each dot represents one donor, corresponding to one independent experimental batch (n = 3). Data are shown as mean ± SD. Statistical significance was assessed using one-way ANOVA followed by Dunnett’s multiple comparisons test versus the control condition. Exact P values are indicated above the bars.

Given the absence of RNA bias at equal dose, this pattern points to a mechanistic hypothesis: Class II genes operate at a lower activation threshold than Class I. To directly test signaling thresholds, intracellular RNA input must be varied while keeping delivery constant. This is possible with DOTAP-complexed RNA. In these tests, Class I required ≥ 0.5 µg/ml for robust induction, whereas 0.1 µg/ml was sufficient for significant Class II activation (Fig. 6b). These data support a lower activation threshold for Class II genes, helping explain why *M. smithii* can still mount strong Class II responses despite reduced uptake. To ask whether this threshold-like separation can be reproduced by varying the strength of TLR8 stimulation with a defined agonist, we performed a dose titration of the synthetic TLR8 agonist TL8-506 (0.001-1 µg/mL) in primary monocytes and quantified representative Class I and Class II transcripts by RT-qPCR. Across this range, TL8-506 induced both programs without a clear low-dose Class II bias (Supplementary Fig. 10).

Overall, these results show that RNA from *M. stadtmanae* and *M. smithii* has comparable intrinsic activity, while Class II genes respond to lower levels of archaeal RNA-TLR8 input than Class I genes, a pattern not recapitulated by titration of a small-molecule TLR8 agonist.

### Differential binding dynamics of NF-**κ**B p65 versus STAT1/2 at Class I and Class II gene promoters

The observation that Class II genes are robustly induced by low concentrations of archaeal RNA, whereas Class I requires higher doses, indicates distinct input sensitivities of the two modules. In whole-cell settings, lower early cell uptake of *M. smithii* effectively mimics a low-input condition, aligning with comparable Class II but weaker Class I responses; conversely, *M. stadtmanae* behaves as a higher-input condition. What might account for this disparity in input sensitivity? Similar patterns have been described for other immune genes, where promoter-specific features, including chromatin accessibility, enhancer context, and transcription-factor input, shape activation thresholds and dose-response behavior^91^. To begin exploring the transcriptional regulation underlying this divergence, we first examined the role of NF-κB p65, a key transcription factor downstream of TLR8 signaling. p65 plays a central role in orchestrating innate immune gene expression and has been implicated in the regulation of many genes upregulated in response to *M. stadtmanae* or *M. smithii* stimulation, including genes from both Class I and Class II categories^92–96^. Our previous studies have shown that *M. stadtmanae* stimulation triggers rapid p65 nuclear translocation, supporting its involvement in archaea-induced signaling^51^. To assess p65 activation dynamics specifically in PBMC-derived monocytes, we stimulated cells with *M. stadtmanae* or *M. smithii* for 1, 2, or 4 hours and evaluated p65 nuclear translocation by confocal microscopy. Nuclear localization of p65 was already detectable at 1 h following *M. stadtmanae* stimulation, whereas *M. smithii* did not induce visible nuclear accumulation until 2 h. Notably, at each time point examined, *M. stadtmanae* stimulation led to significantly higher levels of nuclear p65 than *M. smithii* stimulation (Fig. 7a). This earlier/higher early-stage nuclear p65 signal parallels the stronger early induction of Class I genes under *M. stadtmanae* and prompted us to test whether early p65 engagement contributes to Class I transcription by ChIP-qPCR. To assess early-stage regulation, we performed ChIP-qPCR at 3 h, an interval chosen because transcription-factor binding typically precedes detectable mRNA changes, targeting promoters of representative Class I (*IL6, TNF*) and Class II (*CCL2, CCL8*) genes. *M. stadtmanae* significantly increased p65 occupancy at Class I promoters versus both the unstimulated control and *M. smithii*, whereas *M. smithii* showed no significant change at these loci (Fig. 7b). In contrast, neither stimulus produced significant p65 enrichment at Class II promoters (Fig. 7b). These data support an early, stimulus-dependent engagement of p65 at Class I loci, while Class II induction proceeds via a different regulatory axis.

**Fig. 7.**
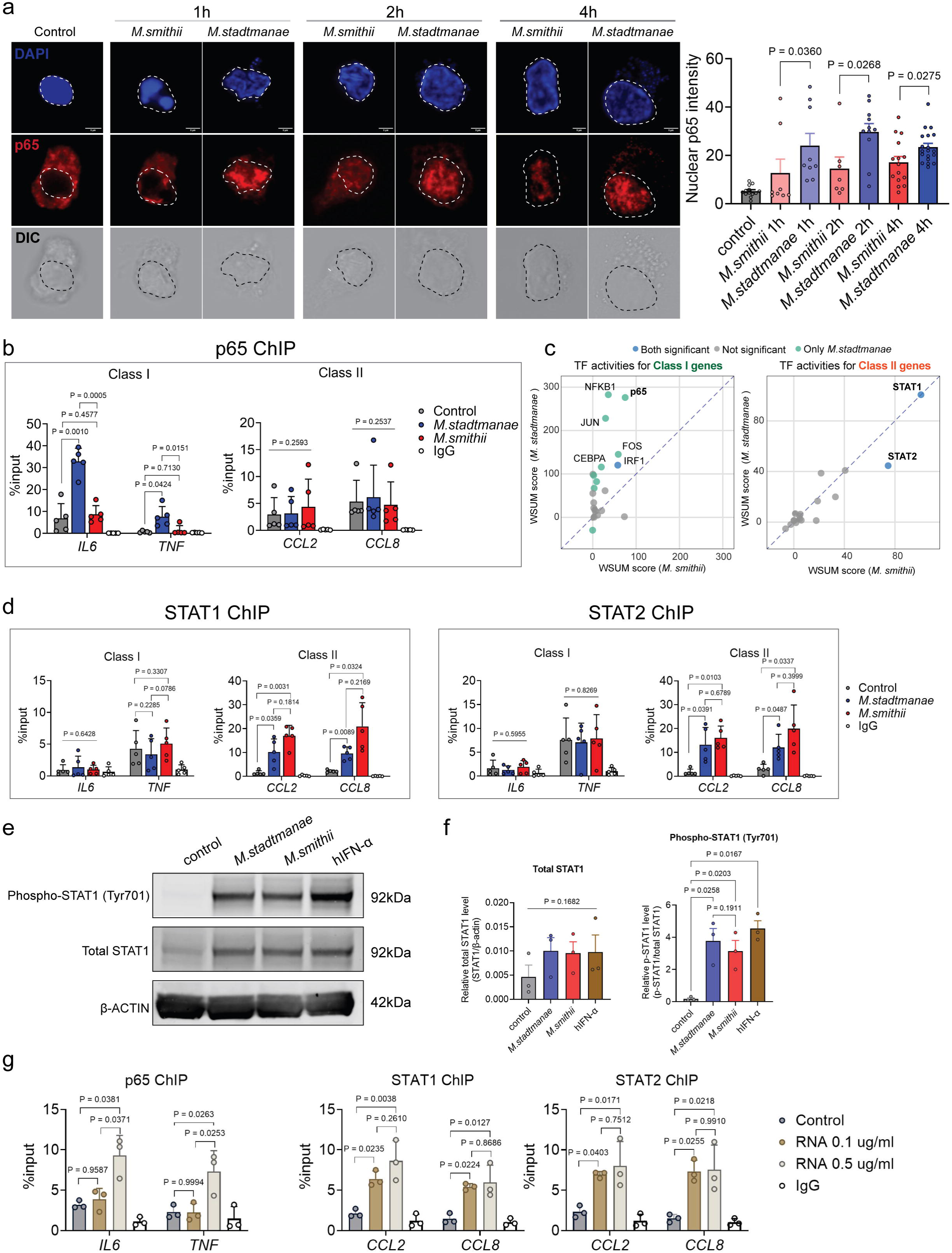
Differential binding dynamics of NF-κB p65 versus STAT1/2 at Class I and Class II gene promoters. **a** Representative confocal immunofluorescence imaging showing nuclear translocation of NF-κB p65 in PBMC-derived monocytes after stimulation with *M. stadtmanae* or *M. smithii* for 1, 2, or 4 h. Nuclear regions were identified using DAPI staining, outlined with dashed circles, and p65 fluorescence intensity within the nuclei was quantified, scale bar = 5 µm. The right panel shows quantification of nuclear fluorescence intensity; each dot represents one cell; 7-19 cells per condition were quantified across four independent donors/batches. Data are shown as mean ± SD. Statistical significance was assessed using the Mann-Whitney test. Exact P values are indicated above the connecting lines. Scale bar = 5 μm. **b** ChIP-qPCR analysis of p65 binding to the promoter regions of selected Class I and Class II genes in PBMC-derived monocytes after 3-h stimulation with *M. stadtmanae* or *M. smithii*. Each dot represents one donor, corresponding to one independent experimental batch (n = 5). Data are shown as percentage of input (% input) and presented as mean ± SD. Statistical significance was determined using one-way ANOVA followed by Tukey’s multiple comparisons test. Exact P values are indicated above the connecting lines. **c** TF activity inference (WSUM) for Class II genes under *M. stadtmanae* and *M. smithii*. The scatter plot shows each transcription factor (one dot per TF). Dot colors encode significance after multiple-testing correction (p < 0.05): blue, significant in both archaea; green, significant only in *M. stadtmanae*; gray, not significant in either stimulus. **d** ChIP-qPCR analysis of STAT1 and STAT2 binding to promoter regions of Class I and Class II genes. PBMC-derived monocytes were stimulated for 3 h with *M. stadtmanae* or *M. smithii*, followed by ChIP-qPCR to assess STAT1 (left) and STAT2 (right) binding to the promoter regions of selected Class I and Class II genes. Each dot represents one donor, corresponding to one independent experimental batch (n = 5). Data are shown as percentage of input (% input) and presented as mean ± SD. Statistical significance was determined using one-way ANOVA followed by Tukey’s multiple comparisons test. Exact P values are indicated above the connecting lines. **e** Representative cropped Western blots showing phospho-STAT1 (Tyr701), total STAT1, and β-actin in primary PBMCs-derived monocytes stimulated with *M. stadtmanae* or *M. smithii* for 3 h, or with human IFN-α for 15 min as a positive control. Molecular weights are indicated as follows: phospho-STAT1 and total STAT1, 92 kDa; β-actin, 42 kDa. Uncropped blots are provided in Supplementary Fig. 13. **f** Quantification of Western blot signals. Left, total STAT1 normalized to β-actin. Right, phospho-STAT1 normalized to total STAT1. Each dot represents one donor, corresponding to one independent experimental batch (n = 3). Data are shown as mean ± SEM. Statistical significance was assessed by one-way ANOVA followed by Tukey’s multiple comparisons test. Exact P values are indicated in the graph. **g** ChIP-qPCR analysis p65, STAT1, and STAT2 binding to promoter regions of selected Class I and Class II genes at different *M. smithii* RNA input concentrations. PBMC-derived monocytes were stimulated for 3 h with purified *M. smithii* RNA at low (0.1 µg/mL) or high (0.5 µg/mL) concentrations. ChIP-qPCR was then performed to assess p65 binding to the promoter regions of selected Class I and Class II genes. Each dot represents one donor, corresponding to one independent experimental batch (n = 3). Data are presented as percentage of input (% input) and shown as mean ± SD. Statistical significance was determined using one-way ANOVA followed by Tukey’s multiple comparisons test. Exact P values are indicated above the connecting lines.

To identify upstream regulators, especially for Class II genes, we performed DoRothEA-based TF enrichment and inferred TF activity using WSUM. As expected, Class I analysis identified 29 TFs, with stronger enrichment under *M. stadtmanae* than *M. smithii* (Fig. 7c). NF-κB subunits ranked highest under *M. stadtmanae*, with NFKB1/p50 and p65 showing higher activity than under *M. smithii*, consistent with increased p65 occupancy at Class I promoters by ChIP-qPCR. For Class II genes, STAT1 and STAT2 were the only TFs significantly enriched under both archaeal stimuli, with comparable activity scores, supporting a shared STAT1/2-centered regulatory axis rather than stimulus-specific TF usage. Full TF lists, activity scores, and statistics are provided in Supplementary Data 4.

STAT1 and STAT2 are central transcription factors of the type I interferon signaling pathway, mediating downstream responses through binding to interferon-stimulated response elements (ISREs) located in the promoter or enhancer regions of target genes^97–99^. Representative Class II genes such as *CCL2*, *CCL8*, and *CXCL10* are well-established STAT1/2 targets^100, 101^. Consistently, HOMER motif analysis of Class II gene regions identified significant enrichment of interferon-related motifs, including IRF1, ISRE, IRF2, IRF8, and IRF3 (adjusted P < 0.01; Supplementary Fig. 11a). Together, these results support an interferon-responsive architecture consistent with STAT1/STAT2-associated Class II regulation. In line with these findings, ChIP-qPCR also showed significant STAT1 and STAT2 binding at Class II promoters (*CCL2*, *CCL8*) upon stimulation with both *M. stadtmanae* and *M. smithii*, with a numerically higher occupancy under *M. smithii* that did not reach significance. Occupancy at the *IL6* and *TNF* promoters remained unchanged (Fig. 7d). These data identify STAT1 and STAT2 as direct regulators of Class II genes, in contrast to the NF-κB-dependent Class I module.

To further assess the STAT1-associated branch, we examined STAT1 phosphorylation and related transcriptional responses after archaeal stimulation. In primary human PBMC-derived monocytes, both *M. stadtmanae* and *M. smithii* induced STAT1 Tyr701 phosphorylation after 3 h, with no detectable difference between species (Fig. 7e, f; Supplementary Fig. 12). IFN-α stimulation for 15 min served as a positive control. Both archaea also modestly increased total STAT1 protein relative to β-actin, consistent with RNA-seq data from PBMCs of nine donors showing significant upregulation of STAT1 and STAT2 at 4 h and 24 h after stimulation with either species (Supplementary Fig. 11b). In the same dataset, IFNB1 was significantly induced at 4 h by both archaea but declined by 24 h. Together, these findings support a shared interferon-responsive program associated with early IFNB1 induction, STAT1 activation, and increased STAT1/2 abundance under both archaeal stimuli.

To directly test whether the differential activation thresholds seen at the input-dose level translate into promoter occupancy, we performed ChIP-qPCR after DOTAP-mediated delivery of purified archaeal RNA at low (0.1 µg/ml) and high (0.5 µg/ml) doses. Because our RNA-delivery experiments showed no intrinsic difference between *M. stadtmanae* and *M. smithii* RNA, we used purified *M. smithii* RNA for the ChIP-qPCR dose test. These concentrations were chosen based on our prior finding that Class II genes can already be induced at 0.1 µg/ml RNA, whereas Class I genes require higher doses (Fig. 6c). Consistent with this, p65 binding to *IL6* and *TNF* promoters was significantly enhanced only at the higher RNA concentration, whereas STAT1 and STAT2 binding to *CCL2* and *CCL8* promoters was already significantly enriched at 0.1 µg/mL and remained similarly high at 0.5 µg/mL (Fig. 7g). Together, these results mirror the transcriptional data and support a model in which, during the early stage of stimulation, Class I genes are preferentially driven by p65 and require stronger archaeal inputs, whereas Class II genes are sustained by a STAT1/2-centered axis that is already engaged at low inputs and remains largely similar across stimuli under equalized RNA, accounting for their consistent induction by both archaea.

## Discussion

Our data indicate that human sensing of gut methanogens resolves, early in stimulation, into two transcriptional programs with distinct input requirements (Fig. 8). We used a stepwise strategy to define the response architecture induced by methanogenic archaea in primary human cells (Supplementary Fig. 13). PBMCs served as the initial donor-matched discovery system, allowing us to capture the overall time-resolved response pattern in a mixed leukocyte setting and to identify the dominant responding compartment. Based on these findings, the mechanistic part of the study was carried out mainly in purified CD14^+^ monocytes, in which we examined uptake, RNA-dependent activation, and the signaling basis of Class I and Class II gene induction. moDCs were included in selected experiments as an additional mononuclear phagocyte population. Recent work further suggests that archaeal host interactions may extend beyond mononuclear phagocytes, as archaeal extracellular vesicles were also shown to interact with intestinal epithelial cells^57^. Against this cellular framework, our data support a model in which Class I genes are preferentially linked to NF-κB p65 and require higher signaling input, whereas Class II genes are associated with a STAT1/2-centered axis that is engaged at lower input levels. The two response modules identified here, may represent distinct layers of TLR8-driven activation with different signaling thresholds.

**Fig. 8.** Proposed model of archaeal uptake and TLR8 signaling in PBMC-derived monocytes. *M. stadtmanae* (left) is taken up more efficiently than *M. smithii* (right), resulting in stronger delivery of archaeal RNA to endosomal TLR8. This stronger input is associated with more pronounced NF-κB (p65) activation together with STAT1/2 associated signaling, inducing both Class I and Class II gene modules. In contrast, the weaker uptake of *M. smithii* is proposed to reduce overall TLR8 engagement and p65-dependent activation, leading to reduced induction of Class I genes. However, Class II genes, which require a lower activation threshold and are associated with STAT1/2, remain largely preserved. Both methanogens induced STAT1 phosphorylation, consistent with engagement of the STAT1/2-associated arm of the response under both conditions. Together, this model suggests that the two gene modules differ in their signaling requirements, with Class I genes being more sensitive to reduced TLR8 input than Class II genes.

Prior studies show that phagocytosis itself can enhance TLR-NF-κB signaling, for instance, uptake of inert beads increases LPS-induced NF-κB activation in macrophage-like cell lines such as U937 and THP-1^102^. This offers a plausible explanation for the stronger early Class I bias observed with *M. stadtmanae*, which exhibits higher early association with phagocytes. That said, directly testing Class I’s dependence on uptake in primary PBMCs is challenging, as blocking uptake tends to abolish multiple signaling inputs simultaneously, making clear mechanistic dissection difficult. In contrast, Class II expression appeared less sensitive to differences in uptake efficiency. The early kinetic data support a model in which relatively limited delivery is already sufficient to trigger this module, particularly for *M. stadtmanae*, whereas *M. smithii* reaches similar expression levels only after further accumulation over time. Notably, once this response is engaged, additional uptake does not translate into proportionally higher Class II output, suggesting that its magnitude may become constrained by regulatory mechanisms such as early negative feedback^103, 104^, receptor desensitization, or cofactor limitation, rather than by further increases in uptake. Future work will address the basis of differential uptake from both the archaeal and host perspectives, with the goal of identifying archaeal features and phagocyte receptors that control recognition and internalization.

TLR8 inhibition almost completely abolished *M. smithii*-induced genes, whereas *M. stadtmanae* left a small residual program. Importantly, the CU-CPT9a-insensitive genes did not resemble a random residual set but instead resolved into coherent transcriptional modules.

The cluster centered on CD276 and IL3RA may point to a distinct myeloid response state linked to immune regulation and cellular interaction^105^, which is particularly interesting in light of the stronger uptake of *M. stadtmanae*. In parallel, the redox-related cluster centered on the SLC7A11-GCLM axis, together with the enrichment of NFE2L2-associated antioxidant and detoxification pathways, indicates a distinct adaptive program that may reflect redox control during archaeal uptake or intracellular processing. The presence of an additional DUSP1-containing immediate-early module further supports the view that these genes represent biologically organized response programs rather than a nonspecific residue after TLR8 inhibition. By contrast, the induction of overt immune response modules remained largely TLR8-dependent. This distinguishes archaeal sensing from bacterial stimulation, where multiple PRRs are often engaged in parallel to drive broader immune activation. This relative simplicity may offer translational advantages over multi-PRR bacterial stimuli. In line with this, clinical-stage TLR8 agonists have shown immunostimulatory activity in antiviral and oncology settings^54, 55^.

DOTAP-RNA delivery experiments suggest that whole-cell differences primarily reflect uptake rather than intrinsic RNA immunostimulatory capacity. The result aligns with established principles: uridine-rich, minimally modified RNAs potently engage TLR7/8 and drive TNF-α/IL-1β^106, 107^. whereas RNAs enriched for modified nucleosides (m C, m A, m U, s²U, pseudouridine) blunt TLR recognition^108^, suggesting no major modification differences between the two archaeal RNAs in our setting. Importantly, these RNA dose-response experiments revealed a threshold-like pattern, with Class II outputs responding at lower RNA doses than Class I. Previous study indicate that human TLR8 senses RNA degradation products through two ligand-binding pockets, with uridine engaging site 1 and a short oligoribonucleotide fragment engaging site 2^109^. In this setting, effective receptor activation depends not only on ligand binding, but also on endolysosomal RNA processing that generates both components, including RNase T2- and RNase 2-dependent steps^110^. Consistent with the importance of this processing layer, pseudouridine-containing RNA was recently shown to evade immune detection because impaired endolysosomal processing limits productive TLR engagement^111^. By contrast, small-molecule TLR8 agonists such as TL8-506 bypass complex RNA degradation and are described as primarily engaging the site 1 uridine pocket^110^. In our titration experiments, TL8-506 did not reproduce a clear low-dose Class II bias. Although our data do not establish the underlying mechanism directly, they are consistent with the idea that the threshold behavior observed with archaeal RNA is shaped by the combined effects of RNA processing and dual-pocket ligand generation, features that may not be fully recapitulated by a site 1-biased small-molecule agonist.

Class I and II were defined operationally by their response patterns rather than by fixed pathway identities. Multiple lines of evidence support a model in which Class I genes are biased toward NF-κB/p65, whereas Class II genes align more strongly with a STAT1/2-centered axis. Whether this generalizes across each class remains to be established. In aggregate, Class I genes (e.g., *IL6*, *TNF*) typically require robust stimulation and show graded, dose-dependent profiles^112, 113^. *IL6* in particular depends on p65 and synergizes with NF-IL6/C/EBPβ^114, 115^, consistent with early high-amplitude but self-limited dynamics. This notion aligns with genome-wide evidence that p65 binding is pervasive but only weakly predictive of gene induction, with many sites are occupied, yet few linked to induced transcripts^116^, whereas STAT1/2 binding under IFN- γ correlates more directly with transcriptional up-regulation^117^. Thus, p65-driven Class I genes likely require higher-order/stronger occupancy to cross an activation threshold, while STAT1/2-driven Class II genes can be engaged under lower inputs. By contrast, the Class II response is more consistent with a STAT1/2-centered regulatory mode. The early induction of *IFNB1* by both *M. stadtmanae* and *M. smithii*, together with the observed increase in total STAT1 protein, the STAT1 phosphorylation pattern, and the broader STAT1/2-associated features of this gene class, supports a model in which relatively modest early input can engage the type I interferon branch and establish a self-reinforcing STAT-centered circuit that helps sustain Class II expression.^97, 99, 118^. Additional mechanisms may facilitate lower effective thresholds for Class II, including chromatin priming (e.g., IFN-γ and LPS synergistically induce *CCL2* via promoter H3K27 acetylation^119^), contexts in which TLR8 can promote IFN-independent STAT1 phosphorylation^120^, and disease-linked hormonal/miRNA pathways that could further enhance STAT signaling and lower thresholds^121^. More broadly, whether this thresholded Class I/II separation is specific to archaeal uptake or also applies to other RNA-based TLR8 inputs is worth exploring. Our dose-response data suggest that careful control of TLR8 input could in principle bias toward Class II at lower doses and recruit Class I only at higher doses, offering a basis to tune cytokine profiles for different applications. However, it should also be noted that the evidence for STAT1/2 involvement is restricted to two representative loci (*CCL2*, *CCL8)* plus increases in STAT1/2 transcripts. In summary we propose a threshold-based model in which Class I genes require strong, uptake-coupled TLR8 input, whereas lower-threshold Class II is compatible with STAT1/2 engagement across a range of inputs. This separation suggests practical means to tune inflammation, for example by adjusting uptake/TLR8 activity or influencing the STAT1/2 axis and motivates biomarker strategies based on archaeal load or uptake features.

## Data availability

The sequencing data are available at European Genome Phenome Archive (EGA) under study accession ID EGAS50000001758. The key results of the RNA-Seq analyses (tables, plots, and interactive HTML QC and results reports) and the Quarto document used for performing transcriptome profiling and for generating respective manuscript figures are available at FAIRDOMHub under https://fairdomhub.org/projects/496. Source data are provided with this paper.

## Author contributions

F.X. and H.H. designed the experiments. F.X. performed the majority of the experiments. J.S. performed multiplex immunoassay experiments. P.Z. performed the RT-qPCR experiments. S.S. performed the western blot experiments. I.W. performed RNA-Seq data analyses. F.X. and H.H. investigated RNA-Seq results with the support of I.W.. F.X. contributed to RNA-Seq figure generation. Cultivation of the methanogenic archaea *M. smithii* and *M. stadtmanae* was performed by V.W. and C.M.E.. F.X. and H.H. wrote the manuscript, H.H. provided funding for the project and conceptional suggestions for its execution.

## Declaration of interests

The authors declare no competing interests.

## Acknowledgements

This work utilized the core infrastructures at the Research Center Borstel, including high-performance computing and fluorescence cytometry. This research was partially funded by the German Research Foundation (DFG, grant HE 2758/6-1, H.H. and S.S.) and the Austrian Science Fund (FWF) [10.55776/F8300], within the framework of the SFB “Immunometabolism” (C.M.E., V.W.).

## Notes

https://fairdomhub.org/projects/496.

### Competing Interest Statement

The authors have declared no competing interest.

### Summary of Updates

In the revised version, we focused first on the central mechanistic point highlighted in your decision letter. We clarified how TLR8 signaling contributes to the differential responses induced by the two archaea, as requested by all reviewers. We strengthened this conclusion by adding experiments in a TLR8-deficient cellular system, which allowed us to distinguish TLR8-dependent response programs from TLR8-independent responses. We performed additional experiments on phagocytosis differences between the two archaea, including staining with archaeon-specific antibodies to validate differential uptake and bead-based phagocytosis assays to assess the contribution of culture. We extended the biochemical analysis of signaling pathways and found that both archaea induce marked STAT1 phosphorylation, supporting a functional role for STAT1 in Class II gene regulation. We also used multiple control experiments to address concerns regarding archaeal stability, potential contaminating factors in the RNA preparation, and the interpretation of possible RNA modification-related differences. All reviewer comments were further addressed individually in the accompanying point-by-point response.

## Reference

1. Oren, A. Molecular ecology of extremely halophilic Archaea and Bacteria. FEMS Microbiol Ecol 39, 1–7 (2002).

2. Kashefi, K. & Lovley, D.R. Extending the upper temperature limit for life. Science 301, 934 (2003).

3. Brochier-Armanet, C., Boussau, B., Gribaldo, S. & Forterre, P. Mesophilic Crenarchaeota: proposal for a third archaeal phylum, the Thaumarchaeota. Nat Rev Microbiol 6, 245–252 (2008).

4. DeLong, E.F. Archaea in coastal marine environments. Proc Natl Acad Sci U S A 89, 5685–5689 (1992).

5. DeLong, E.F. Everything in moderation: archaea as ’non-extremophiles’. Curr Opin Genet Dev 8, 649–654 (1998).

6. Moissl-Eichinger, C. et al. Human age and skin physiology shape diversity and abundance of Archaea on skin. Sci Rep 7, 4039 (2017).

7. Pausan, M.R. et al. Exploring the Archaeome: Detection of Archaeal Signatures in the Human Body. Front Microbiol 10, 2796 (2019).

8. Koskinen, K. et al. First Insights into the Diverse Human Archaeome: Specific Detection of Archaea in the Gastrointestinal Tract, Lung, and Nose and on Skin. mBio 8 (2017).

9. Li, C.L. et al. Prevalence and molecular diversity of Archaea in subgingival pockets of periodontitis patients. Oral Microbiol Immunol 24, 343–346 (2009).

10. Lepp, P.W. et al. Methanogenic Archaea and human periodontal disease. Proc Natl Acad Sci U S A 101, 6176–6181 (2004).

11. Kulik, E.M., Sandmeier, H., Hinni, K. & Meyer, J. Identification of archaeal rDNA from subgingival dental plaque by PCR amplification and sequence analysis. FEMS Microbiol Lett 196, 129–133 (2001).

12. Horz, H.P. & Conrads, G. Methanogenic Archaea and oral infections - ways to unravel the black box. J Oral Microbiol 3 (2011).

13. Sogodogo, E. et al. Nine Cases of Methanogenic Archaea in Refractory Sinusitis, an Emerging Clinical Entity. Front Public Health 7, 38 (2019).

14. Baehren, C. et al. The Relevance of the Bacterial Microbiome, Archaeome and Mycobiome in Pediatric Asthma and Respiratory Disorders. Cells 11 (2022).

15. Blais Lecours, P., et al. Increased Prevalence of Methanosphaera stadtmanae in Inflammatory Bowel Diseases. PLoS One 9, e87734 (2014).

16. Candeliere, F., Sola, L., Raimondi, S., Rossi, M. & Amaretti, A. Good and bad dispositions between archaea and bacteria in the human gut: New insights from metagenomic survey and co-occurrence analysis. Synth Syst Biotechnol 9, 88–98 (2024).

17. Chibani, C.M. et al. A catalogue of 1,167 genomes from the human gut archaeome. Nat Microbiol 7, 48–61 (2022).

18. Dridi, B., Henry, M., El Khechine, A., Raoult, D. & Drancourt, M. High prevalence of Methanobrevibacter smithii and Methanosphaera stadtmanae detected in the human gut using an improved DNA detection protocol. PLoS One 4, e7063 (2009).

19. Mbakwa, C.A. et al. Gut colonization with methanobrevibacter smithii is associated with childhood weight development. Obesity (Silver Spring) 23, 2508–2516 (2015).

20. Bai, X. et al. Landscape of the gut archaeome in association with geography, ethnicity, urbanization, and diet in the Chinese population. Microbiome 10, 147 (2022).

21. Ruaud, A. et al. Syntrophy via Interspecies H(2) Transfer between Christensenella and Methanobrevibacter Underlies Their Global Cooccurrence in the Human Gut. mBio 11 (2020).

22. Pasalari, H., Gholami, M., Rezaee, A., Esrafili, A. & Farzadkia, M. Perspectives on microbial community in anaerobic digestion with emphasis on environmental parameters: A systematic review. Chemosphere 270, 128618 (2021).

23. Dirks, B. et al. Methanogenesis associated with altered microbial production of short-chain fatty acids and human-host metabolizable energy. ISME J 19 (2025).

24. Li, T. et al. Multi-Cohort Analysis Reveals Altered Archaea in Colorectal Cancer Fecal Samples Across Populations. Gastroenterology 168, 525–538 e522 (2025).

25. Klindworth, A. et al. Evaluation of general 16S ribosomal RNA gene PCR primers for classical and next-generation sequencing-based diversity studies. Nucleic Acids Res 41, e1 (2013).

26. Low, A. et al. Mutual Exclusion of Methanobrevibacter Species in the Human Gut Microbiota Facilitates Directed Cultivation of a Candidatus Methanobrevibacter Intestini Representative. Microbiol Spectr 10, e0084922 (2022).

27. Weinberger, V. et al. Expanding the cultivable human archaeome: Methanobrevibacter intestini sp. nov. and strain Methanobrevibacter smithii ’GRAZ-2’ from human faeces. Int J Syst Evol Microbiol **75** (2025).

28. Mohammadzadeh, R., Mahnert, A., Duller, S. & Moissl-Eichinger, C. Archaeal key-residents within the human microbiome: characteristics, interactions and involvement in health and disease. Curr Opin Microbiol 67, 102146 (2022).

29. Mohammadzadeh, R. et al. Age-related dynamics of predominant methanogenic archaea in the human gut microbiome. BMC Microbiol 25, 193 (2025).

30. Wang, T. et al. Methanogen Levels Are Significantly Associated with Fecal Microbiota Composition and Alpha Diversity in Healthy Adults and Irritable Bowel Syndrome Patients. Microbiol Spectr 10, e0165322 (2022).

31. Chen, S. et al. Consistent signatures in the human gut microbiome of longevous populations. Gut Microbes 16, 2393756 (2024).

32. Ghoshal, U., Shukla, R., Srivastava, D. & Ghoshal, U.C. Irritable Bowel Syndrome, Particularly the Constipation-Predominant Form, Involves an Increase in Methanobrevibacter smithii, Which Is Associated with Higher Methane Production. Gut Liver 10, 932–938 (2016).

33. Wallen, Z.D. et al. Metagenomics of Parkinson’s disease implicates the gut microbiome in multiple disease mechanisms. Nat Commun 13, 6958 (2022).

34. Mohammadzadeh, R. et al. Methanobrevibacter smithii associates with colorectal cancer through trophic control of the cancer bacteriome. *bioRxiv*, 2025.2006.2012.659283 (2025).

35. Ghavami, S.B. et al. Alterations of the human gut Methanobrevibacter smithii as a biomarker for inflammatory bowel diseases. Microb Pathog 117, 285–289 (2018).

36. Hoegenauer, C., Hammer, H.F., Mahnert, A. & Moissl-Eichinger, C. Methanogenic archaea in the human gastrointestinal tract. Nat Rev Gastroenterol Hepatol 19, 805–813 (2022).

37. Barnett, D.J.M., Mommers, M., Penders, J., Arts, I.C.W. & Thijs, C. Intestinal archaea inversely associated with childhood asthma. J Allergy Clin Immunol 143, 2305–2307 (2019).

38. Koh, A., De Vadder, F., Kovatcheva-Datchary, P. & Backhed, F. From Dietary Fiber to Host Physiology: Short-Chain Fatty Acids as Key Bacterial Metabolites. Cell 165, 1332–1345 (2016).

39. Palm, N.W., de Zoete, M.R. & Flavell, R.A. Immune-microbiota interactions in health and disease. Clin Immunol 159, 122–127 (2015).

40. Round, J.L. & Mazmanian, S.K. Inducible Foxp3+ regulatory T-cell development by a commensal bacterium of the intestinal microbiota. Proc Natl Acad Sci U S A 107, 12204–12209 (2010).

41. Levy, M. et al. Microbiota-Modulated Metabolites Shape the Intestinal Microenvironment by Regulating NLRP6 Inflammasome Signaling. Cell 163, 1428–1443 (2015).

42. Fitzgerald, K.A. & Kagan, J.C. Toll-like Receptors and the Control of Immunity. Cell 180, 1044–1066 (2020).

43. Caruso, R., Warner, N., Inohara, N. & Nunez, G. NOD1 and NOD2: signaling, host defense, and inflammatory disease. Immunity 41, 898–908 (2014).

44. Dalpke, A., Frank, J., Peter, M. & Heeg, K. Activation of toll-like receptor 9 by DNA from different bacterial species. Infection and immunity 74, 940–946 (2006).

45. Eigenbrod, T., Pelka, K., Latz, E., Kreikemeyer, B. & Dalpke, A.H. TLR8 Senses Bacterial RNA in Human Monocytes and Plays a Nonredundant Role for Recognition of Streptococcus pyogenes. J Immunol 195, 1092–1099 (2015).

46. Bieback, K. et al. Hemagglutinin protein of wild-type measles virus activates toll-like receptor 2 signaling. J Virol 76, 8729–8736 (2002).

47. Shimizu, T. RNA recognition in toll-like receptor signaling. Curr Opin Struct Biol 88, 102913 (2024).

48. Kuehnast, T. et al. Exploring the human archaeome: its relevance for health and disease, and its complex interplay with the human immune system. FEBS J 292, 1316–1329 (2025).

49. Bang, C., Weidenbach, K., Gutsmann, T., Heine, H. & Schmitz, A.R. The Intestinal Archaea Methanosphaera stadtmanae and Methanobrevibacter smithii Activate Human Dendritic Cells. PLoS ONE 9, e99411 (2014).

50. Blais Lecours, P., et al. Immunogenic properties of archaeal species found in bioaerosols. PLoS One 6, e23326 (2011).

51. Vierbuchen, T., Bang, C., Rosigkeit, H., Schmitz, R.A. & Heine, H. The Human-Associated Archaeon Methanosphaera stadtmanae Is Recognized through Its RNA and Induces TLR8-Dependent NLRP3 Inflammasome Activation. Front Immunol 8, 1535 (2017).

52. Stein, K. et al. Endosomal recognition of Lactococcus lactis G121 and its RNA by dendritic cells is key to its allergy-protective effects. J Allergy Clin Immunol 139, 667–678 e665 (2017).

53. Zhao, T. et al. Vaccine adjuvants: mechanisms and platforms. Signal Transduct Target Ther 8, 283 (2023).

54. Chow, L.Q.M. et al. Phase Ib Trial of the Toll-like Receptor 8 Agonist, Motolimod (VTX-2337), Combined with Cetuximab in Patients with Recurrent or Metastatic SCCHN. Clin Cancer Res **23**, 2442-2450 (2017).

55. Gane, E.J. et al. Safety and efficacy of the oral TLR8 agonist selgantolimod in individuals with chronic hepatitis B under viral suppression. J Hepatol 78, 513–523 (2023).

56. Oka, S. et al. Archaeal Glycerolipids Are Recognized by C-Type Lectin Receptor Mincle. J Am Chem Soc 145, 18538–18548 (2023).

57. Weinberger, V. et al. Proteomic and metabolomic profiling of extracellular vesicles produced by human gut archaea. Nat Commun 16, 5094 (2025).

58. Fricke, W.F. et al. The genome sequence of Methanosphaera stadtmanae reveals why this human intestinal archaeon is restricted to methanol and H2 for methane formation and ATP synthesis. J Bacteriol 188, 642–658 (2006).

59. Gupta, A.B. & Seedorf, H. Structural and functional insights from the sequences and complex domain architecture of adhesin-like proteins from Methanobrevibacter smithii and Methanosphaera stadtmanae. Front Microbiol 15, 1463715 (2024).

60. Boyum, A. Separation of White Blood Cells. Nature 204, 793–794 (1964).

61. Crifo, B. et al. Hydroxylase Inhibition Selectively Induces Cell Death in Monocytes. J Immunol 202, 1521–1530 (2019).

62. Sallusto, F. & Lanzavecchia, A. Efficient presentation of soluble antigen by cultured human dendritic cells is maintained by granulocyte/macrophage colony-stimulating factor plus interleukin 4 and downregulated by tumor necrosis factor alpha. J Exp Med 179, 1109–1118 (1994).

63. Rapino, F. et al. C/EBPalpha induces highly efficient macrophage transdifferentiation of B lymphoma and leukemia cell lines and impairs their tumorigenicity. Cell Rep 3, 1153–1163 (2013).

64. Maloley, P.M. et al. Performance of a commercially available multiplex platform in the assessment of circulating cytokines and chemokines in patients with rheumatoid arthritis and osteoarthritis. J Immunol Methods 495, 113048 (2021).

65. Bray, N.L., Pimentel, H., Melsted, P. & Pachter, L. Erratum: Near-optimal probabilistic RNA-seq quantification. Nat Biotechnol 34, 888 (2016).

66. Pimentel, H., Bray, N.L., Puente, S., Melsted, P. & Pachter, L. Differential analysis of RNA-seq incorporating quantification uncertainty. Nat Methods 14, 687–690 (2017).

67. Ewels, P., Magnusson, M., Lundin, S. & Kaller, M. MultiQC: summarize analysis results for multiple tools and samples in a single report. Bioinformatics 32, 3047–3048 (2016).

68. Xiao, Y. et al. A novel significance score for gene selection and ranking. Bioinformatics 30, 801–807 (2014).

69. Kanehisa, M., Furumichi, M., Sato, Y., Matsuura, Y. & Ishiguro-Watanabe, M. KEGG: biological systems database as a model of the real world. Nucleic Acids Res 53, D672–D677 (2025).

70. Duller, S. et al. Targeted isolation of Methanobrevibacter strains from fecal samples expands the cultivated human archaeome. Nat Commun 15, 7593 (2024).

71. Kim, T.H. & Dekker, J. ChIP-Quantitative Polymerase Chain Reaction (ChIP-qPCR). Cold Spring Harb Protoc 2018 (2018).

72. Nelson, J.D., Denisenko, O. & Bomsztyk, K. Protocol for the fast chromatin immunoprecipitation (ChIP) method. Nat Protoc 1, 179–185 (2006).

73. Badia, I.M.P., et al. decoupleR: ensemble of computational methods to infer biological activities from omics data. Bioinform Adv 2, vbac016 (2022).

74. Garcia-Alonso, L., Holland, C.H., Ibrahim, M.M., Turei, D. & Saez-Rodriguez, J. Benchmark and integration of resources for the estimation of human transcription factor activities. Genome Res 29, 1363–1375 (2019).

75. Heinz, S. et al. Simple combinations of lineage-determining transcription factors prime cis-regulatory elements required for macrophage and B cell identities. Mol Cell 38, 576–589 (2010).

76. Xu, S. et al. Using clusterProfiler to characterize multiomics data. Nat Protoc 19, 3292–3320 (2024).

77. Kolde, R. (R package version 1.0.13, 2025).

78. Djoko, K.Y., Ong, C.L., Walker, M.J. & McEwan, A.G. The Role of Copper and Zinc Toxicity in Innate Immune Defense against Bacterial Pathogens. J Biol Chem 290, 18954–18961 (2015).

79. Waller, K. et al. ADAM17-Mediated Reduction in CD14(++)CD16(+) Monocytes ex vivo and Reduction in Intermediate Monocytes With Immune Paresis in Acute Pancreatitis and Acute Alcoholic Hepatitis. Front Immunol 10, 1902 (2019).

80. Dale, D.C., Boxer, L. & Liles, W.C. The phagocytes: neutrophils and monocytes. Blood 112, 935–945 (2008).

81. Grinter, R. & Greening, C. Cofactor F420: an expanded view of its distribution, biosynthesis and roles in bacteria and archaea. FEMS Microbiol Rev 45 (2021).

82. Lambrecht, J. et al. Flow cytometric quantification, sorting and sequencing of methanogenic archaea based on F(420) autofluorescence. Microb Cell Fact 16, 180 (2017).

83. Minnebo, Y.D.P., K.; Props, R.; Lacoere, T.; Boon, N.; Van de Wiele, T. Methanogenic Archaea Quantification in the Human Gut Microbiome with F420 Autofluorescence-Based Flow Cytometry. Appl. Microbiol. 4, 162–180 (2024).

84. Zhang, S. et al. Small-molecule inhibition of TLR8 through stabilization of its resting state. Nat Chem Biol 14, 58–64 (2018).

85. Hu, Z. et al. Small-Molecule TLR8 Antagonists via Structure-Based Rational Design. Cell Chem Biol 25, 1286–1291 e1283 (2018).

86. Moen, S.H. et al. Human Toll-like Receptor 8 (TLR8) Is an Important Sensor of Pyogenic Bacteria, and Is Attenuated by Cell Surface TLR Signaling. Front Immunol 10, 1209 (2019).

87. Heil, F. et al. Species-specific recognition of single-stranded RNA via toll-like receptor 7 and 8. Science 303, 1526–1529 (2004).

88. Colak, E. et al. RNA and imidazoquinolines are sensed by distinct TLR7/8 ectodomain sites resulting in functionally disparate signaling events. J Immunol 192, 5963–5973 (2014).

89. Alharbi, A.S. et al. 2’-O-Methyl-guanosine RNA fragments antagonize TLR7 and TLR8 to limit autoimmunity. Nat Immunol 27, 762–775 (2026).

90. Herster, F. et al. Neutrophil extracellular trap-associated RNA and LL37 enable self-amplifying inflammation in psoriasis. Nat Commun 11, 105 (2020).

91. Medzhitov, R. & Horng, T. Transcriptional control of the inflammatory response. Nat Rev Immunol 9, 692–703 (2009).

92. Delerive, P. et al. Peroxisome proliferator-activated receptor alpha negatively regulates the vascular inflammatory gene response by negative cross-talk with transcription factors NF-kappaB and AP-1. J Biol Chem 274, 32048–32054 (1999).

93. Deng, X. et al. Transcriptional regulation of increased CCL2 expression in pulmonary fibrosis involves nuclear factor-kappaB and activator protein-1. Int J Biochem Cell Biol 45, 1366–1376 (2013).

94. Hong, J.H. & Lee, Y.C. Anti-Inflammatory Effects of Cicadidae Periostracum Extract and Oleic Acid through Inhibiting Inflammatory Chemokines Using PCR Arrays in LPS-Induced Lung inflammation In Vitro. Life (Basel) 12 (2022).

95. Shebzukhov Iu, V. & Kuprash, D.V. [Transcriptional regulation of TNF/LT locus in immune cells]. Mol Biol (Mosk) 45, 56–67 (2011).

96. Unger, A. et al. Chromatin Binding of c-REL and p65 Is Not Limiting for Macrophage IL12B Transcription During Immediate Suppression by Ovarian Carcinoma Ascites. Front Immunol 9, 1425 (2018).

97. Platanitis, E. et al. A molecular switch from STAT2-IRF9 to ISGF3 underlies interferon-induced gene transcription. Nat Commun 10, 2921 (2019).

98. Mostafavi, S. et al. Parsing the Interferon Transcriptional Network and Its Disease Associations. Cell 164, 564–578 (2016).

99. Ivashkiv, L.B. & Donlin, L.T. Regulation of type I interferon responses. Nat Rev Immunol 14, 36–49 (2014).

100. Buttmann, M., Berberich-Siebelt, F., Serfling, E. & Rieckmann, P. Interferon-beta is a potent inducer of interferon regulatory factor-1/2-dependent IP-10/CXCL10 expression in primary human endothelial cells. J Vasc Res 44, 51–60 (2007).

101. Blaszczyk, K. et al. STAT2/IRF9 directs a prolonged ISGF3-like transcriptional response and antiviral activity in the absence of STAT1. Biochem J 466, 511–524 (2015).

102. Ueno, T., Yamamoto, Y. & Kawasaki, K. Phagocytosis of microparticles increases responsiveness of macrophage-like cell lines U937 and THP-1 to bacterial lipopolysaccharide and lipopeptide. Sci Rep 11, 6782 (2021).

103. Gilchrist, M. et al. Systems biology approaches identify ATF3 as a negative regulator of Toll-like receptor 4. Nature 441, 173–178 (2006).

104. Boone, D.L. et al. The ubiquitin-modifying enzyme A20 is required for termination of Toll-like receptor responses. Nat Immunol 5, 1052–1060 (2004).

105. Zhang, G. et al. Soluble CD276 (B7-H3) is released from monocytes, dendritic cells and activated T cells and is detectable in normal human serum. Immunology 123, 538–546 (2008).

106. Diebold, S.S., Kaisho, T., Hemmi, H., Akira, S. & Reis e Sousa, C. Innate antiviral responses by means of TLR7-mediated recognition of single-stranded RNA. Science 303, 1529–1531 (2004).

107. Eigenbrod, T. & Dalpke, A.H. Bacterial RNA: An Underestimated Stimulus for Innate Immune Responses. J Immunol 195, 411–418 (2015).

108. Kariko, K., Buckstein, M., Ni, H. & Weissman, D. Suppression of RNA recognition by Toll-like receptors: the impact of nucleoside modification and the evolutionary origin of RNA. Immunity 23, 165–175 (2005).

109. Tanji, H. et al. Toll-like receptor 8 senses degradation products of single-stranded RNA. Nat Struct Mol Biol 22, 109–115 (2015).

110. Greulich, W. et al. TLR8 Is a Sensor of RNase T2 Degradation Products. Cell 179, 1264–1275 e1213 (2019).

111. Berouti, M. et al. Pseudouridine RNA avoids immune detection through impaired endolysosomal processing and TLR engagement. Cell 188, 4880–4895 e4815 (2025).

112. Natoli, G., Saccani, S., Bosisio, D. & Marazzi, I. Interactions of NF-kappaB with chromatin: the art of being at the right place at the right time. Nat Immunol 6, 439–445 (2005).

113. Werner, S.L., Barken, D. & Hoffmann, A. Stimulus specificity of gene expression programs determined by temporal control of IKK activity. Science 309, 1857–1861 (2005).

114. Matsusaka, T. et al. Transcription factors NF-IL6 and NF-kappa B synergistically activate transcription of the inflammatory cytokines, interleukin 6 and interleukin 8. Proc Natl Acad Sci U S A 90, 10193–10197 (1993).

115. Libermann, T.A. & Baltimore, D. Activation of interleukin-6 gene expression through the NF-kappa B transcription factor. Mol Cell Biol 10, 2327–2334 (1990).

116. Jurida, L. et al. The Activation of IL-1-Induced Enhancers Depends on TAK1 Kinase Activity and NF-kappaB p65. Cell Rep 10, 726–739 (2015).

117. Satoh, J. & Tabunoki, H. A Comprehensive Profile of ChIP-Seq-Based STAT1 Target Genes Suggests the Complexity of STAT1-Mediated Gene Regulatory Mechanisms. Gene Regul Syst Bio 7, 41–56 (2013).

118. Michalska, A., Blaszczyk, K., Wesoly, J. & Bluyssen, H.A.R. A Positive Feedback Amplifier Circuit That Regulates Interferon (IFN)-Stimulated Gene Expression and Controls Type I and Type II IFN Responses. Front Immunol 9, 1135 (2018).

119. Akhter, N. et al. IFN-gamma and LPS Induce Synergistic Expression of CCL2 in Monocytic Cells via H3K27 Acetylation. J Inflamm Res 15, 4291–4302 (2022).

120. Larange, A., Antonios, D., Pallardy, M. & Kerdine-Romer, S. TLR7 and TLR8 agonists trigger different signaling pathways for human dendritic cell maturation. J Leukoc Biol 85, 673–683 (2009).

121. Young, N.A. et al. Estrogen-regulated STAT1 activation promotes TLR8 expression to facilitate signaling via microRNA-21 in systemic lupus erythematosus. Clin Immunol 176, 12–22 (2017).

